# Mapping Functional Dynamics Hotspots for Protein Engineering with NMR Peak Intensity Analysis

**DOI:** 10.1101/2025.10.26.684639

**Authors:** Adam M. Damry, Serena E. Hunt, Sandrine Legault, Michael C. Thompson, Natalie K. Goto, Roberto A. Chica

## Abstract

Structural dynamics play a crucial role in protein function, and tuning these dynamics through mutagenesis has emerged as a promising strategy for enhancing activity. However, identifying dynamics hotspots for protein engineering remains a labor-intensive challenge. Here, we demonstrate that NMR peak intensity analysis—a rapid, qualitative method with residue-level resolution—can identify functionally relevant dynamic regions with high precision. Using a family of red fluorescent proteins (RFPs) as a case study, we reveal that flexibility in specific regions of their structures correlates with function. Specifically, as quantum yield increases, the side of the β-barrel closest to the chromophore phenolate moiety becomes more rigid, while the opposite side, closest to the acylimine group, gains flexibility. Notably, the phenolate face corresponds to a mutational hotspot frequently targeted in directed evolution campaigns aimed at enhancing brightness, underscoring its functional significance. B-factor analysis of non-cryogenic X-ray crystal structures further supports our findings. Our results establish NMR peak intensity analysis as a promising tool for mapping functional dynamics hotspots to guide protein engineering campaigns.

## Introduction

Structural dynamics play a critical role in a wide range of protein functions, including ligand binding ^1^, enzyme catalysis ^2^, and allosteric regulation ^3^. Tuning these internal motions through targeted mutations has been shown to enhance function ^4,5^. Yet, identifying which dynamic regions contribute most significantly to function remains a time-consuming and technically demanding task ^6^, limiting this protein engineering pathway to well-characterized systems. Nuclear magnetic resonance (NMR) spectroscopy is uniquely suited for characterizing protein dynamics at atomic resolution under near-physiological conditions, with standard approaches such as relaxation dispersion or Carr-Purcell-Meiboom-Gill (CPMG) experiments ^7,8^ providing quantitative high-resolution dynamical information. However, it is not always immediately clear which amongst many NMR experiments would be most appropriate to measure dynamics in a protein system, as this selection depends on timescales of exchange and populations of excited states ^9–11^. For the purposes of protein engineering, a higher-throughput approach that could identify regions where conformational dynamics correlate with function would be especially valuable for comparing multiple variants.

Building on the established relationship between NMR peak lineshapes and dynamics ^12–14^, we propose peak intensity analysis as a practical yet underutilized approach for probing functional protein dynamics. This approach infers dynamic information from ^1^H–^15^N Heteronuclear Single Quantum Coherence (HSQC) peak intensities, since they are influenced by both fast (ps–ns) ^13^ and intermediate (µs–ms) ^14^ timescale motions. Specifically, micro-to millisecond exchange processes enhance transverse relaxation rates, causing peak broadening that reduces peak intensity. Conversely, pico-to nanosecond fluctuations that are faster than the global molecular tumbling rate suppress relaxation and enhance peak intensity and sharpness. Crucially, a single assigned spectrum is sufficient to begin analysis, as HSQC spectra of closely related mutants are often highly similar, making this technique well-suited to the comparative study of engineered protein variants.

Here, we demonstrate that NMR peak intensity analysis can be used to identify functionally relevant dynamics hotspots. Using a family of red fluorescent proteins (RFPs) as a model system, we apply this approach to link structural flexibility to quantum yield, a property directly influenced by protein dynamics. Our results reveal that as quantum yield increases, the side of the RFP β-barrel closest to the chromophore phenolate moiety becomes more rigid, while the opposite side, closest to the acylimine group, becomes more flexible. Notably, the phenolate face corresponds to a mutational hotspot frequently targeted in directed evolution campaigns aimed at enhancing brightness. We validate these findings using B-factor analysis of non-cryogenic X-ray crystal structures. Together, our results establish NMR peak intensity analysis as a rapid and accessible first pass method for identifying functionally relevant dynamic regions, potentially accelerating the discovery of beneficial mutations in protein engineering.

## Results

### Red fluorescent proteins as a model for function-linked dynamics

To explore whether NMR peak intensity analysis can rapidly identify functionally relevant dynamics hotspots, we investigated a family of RFPs due to the well-established connection between chromophore flexibility and fluorescence quantum yield: rigid chromophore environments enhance radiative decay, while increased motion leads to energy loss through non-radiative pathways ^15–17^. We hypothesized that applying NMR peak intensity analysis to a panel of RFPs spanning a wide range of quantum yields would reveal localized dynamics changes correlating with fluorescence output, offering a rapid approach for identifying functionally critical dynamic regions.

We selected mCherry ^18^, a well-characterized monomeric RFP optimized through mutagenesis for improved brightness and photostability, along with four engineered variants ^19–21^, each containing between 6 and 30 mutations (Figure 1, Supplementary Tables 1–2, Supplementary Figure 1). Together, these five RFPs span a range of quantum yields of 0.02 to 0.75 (Table 1, Supplementary Figure 2), with excitation and emission maxima ranging from 569–604 nm and 592–633 nm, respectively. With molecular weights of approximately 27 kDa (Supplementary Table 1), these proteins represent a tractable system for NMR-based dynamic analysis. Importantly, all mCherry variants share the same chromophore derived from the MYG tripeptide (Supplementary Figure 3), indicating that differences in quantum yield arise from environmental factors, such as structural or dynamic variations, rather than changes in chromophore chemistry. Furthermore, all variants have resolved crystal structures, enabling three-dimensional mapping of peak intensity-derived dynamics.

**Figure 1.**
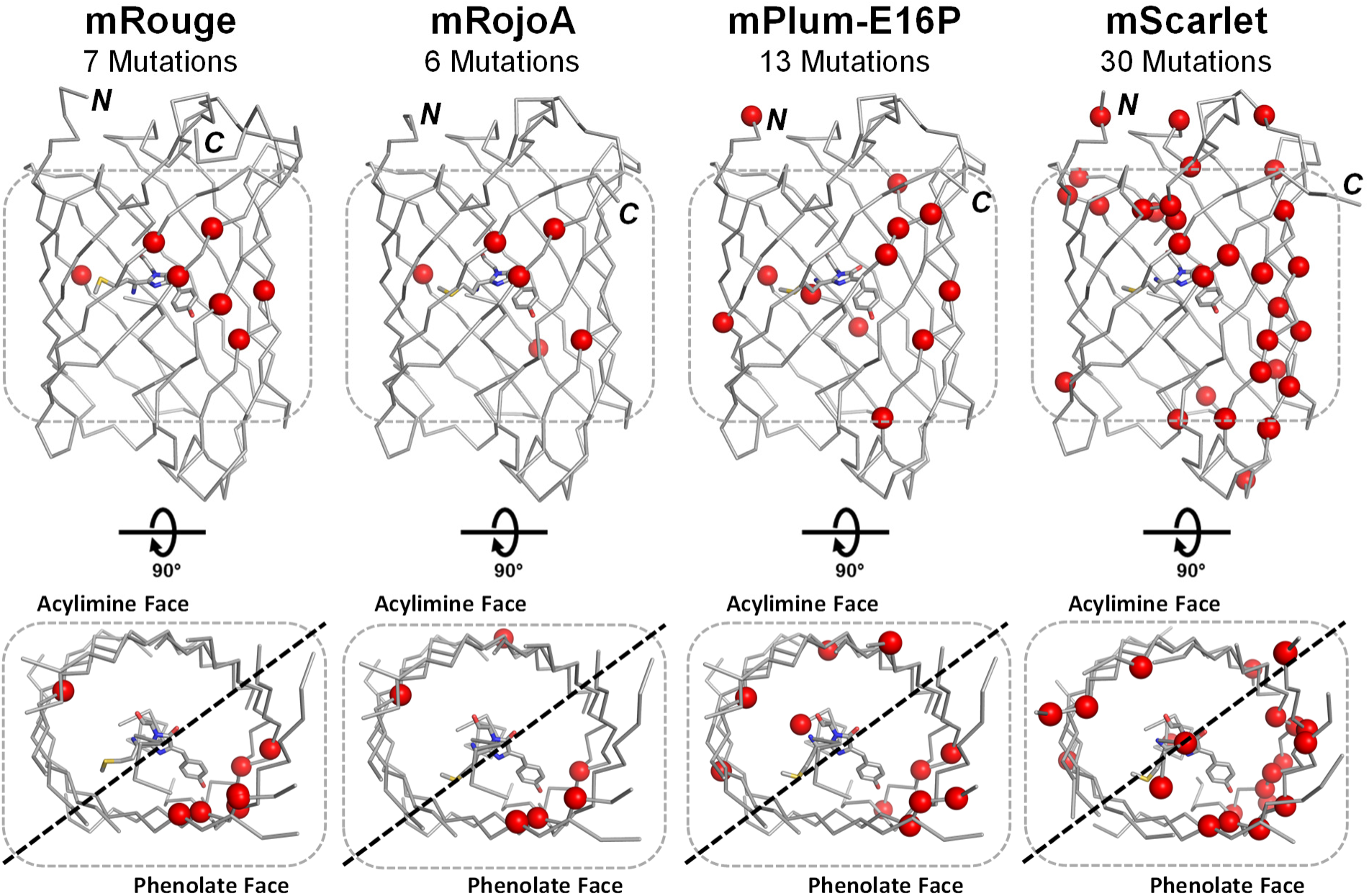
mCherry variants. Crystal structures of monomeric RFPs mRouge (PDB: 3NED) ^20^, mRojoA (PDB: 3NEZ) ^20^, mPlum-E16P (PDB: 4H3L) ^21^, and mScarlet (PDB: 5LK4) ^19^ are shown as ribbon diagrams. Mutations relative to mCherry are shown as spheres, chromophores as sticks, and N- and C-termini are labeled.

**Table 1.**
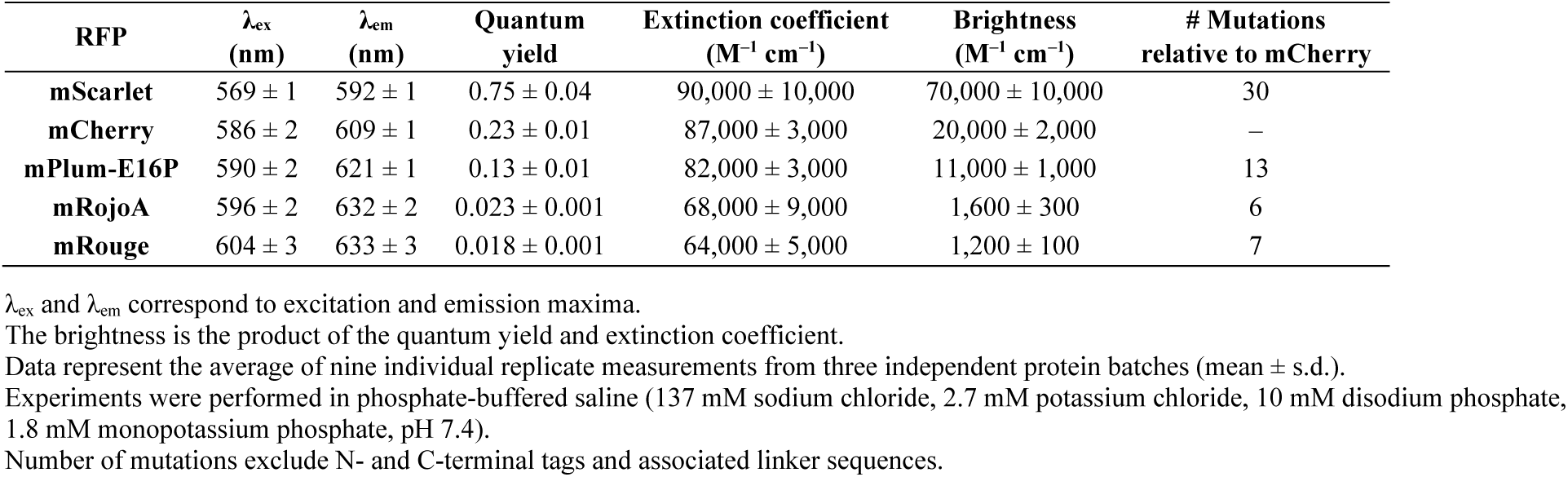
Properties of RFPs.

| RFP | $\lambda_{\text{ex}}$<br>(nm) | $\lambda_{\text{em}}$<br>(nm) | Quantum<br>yield | Extinction coefficient<br>(M<sup>-1</sup> cm<sup>-1</sup>) | Brightness<br>(M<sup>-1</sup> cm<sup>-1</sup>) | # Mutations<br>relative to mCherry |
| --- | --- | --- | --- | --- | --- | --- |
| mScarlet | 569 ± 1 | 592 ± 1 | 0.75 ± 0.04 | 90,000 ± 10,000 | 70,000 ± 10,000 | 30 |
| mCherry | 586 ± 2 | 609 ± 1 | 0.23 ± 0.01 | 87,000 ± 3,000 | 20,000 ± 2,000 | — |
| mPlum-E16P | 590 ± 2 | 621 ± 1 | 0.13 ± 0.01 | 82,000 ± 3,000 | 11,000 ± 1,000 | 13 |
| mRojoA | 596 ± 2 | 632 ± 2 | 0.023 ± 0.001 | 68,000 ± 9,000 | 1,600 ± 300 | 6 |
| mRouge | 604 ± 3 | 633 ± 3 | 0.018 ± 0.001 | 64,000 ± 5,000 | 1,200 ± 100 | 7 |

$\lambda_{\text{ex}}$ and $\lambda_{\text{em}}$ correspond to excitation and emission maxima. The brightness is the product of the quantum yield and extinction coefficient. Data represent the average of nine individual replicate measurements from three independent protein batches (mean ± s.d.). Experiments were performed in phosphate-buffered saline (137 mM sodium chloride, 2.7 mM potassium chloride, 10 mM disodium phosphate, 1.8 mM monopotassium phosphate, pH 7.4). Number of mutations exclude N- and C-terminal tags and associated linker sequences.

### Peak intensity analysis reveals dynamic signatures correlated with quantum yield

We acquired and assigned ^1^H–^15^N HSQC spectra for all five RFPs (Supplementary Figures 4–8). The dimmest RFPs, mRojoA and mRouge, displayed substantial peak broadening (Supplementary Figure 9), which reduced the fraction of assignable residues (69–82%) compared to the brighter variants (≥90%), consistent with increased structural dynamics on intermediate timescales. Notably, mRojoA showed precipitation and fluorescence loss during data acquisition, leading to the lowest assignment coverage among all variants. To improve coverage in the low-brightness regime, we analyzed the spectra of mRojoA and mRouge in combination, exploiting their high sequence identity (98%, six mutations), and achieved 88% coverage across both dim variants, comparable to that obtained for brighter proteins.

To gain insight into the dynamics of these variants at the residue level, we measured intensities of peaks in HSQC spectra for each variant. To identify residues with dynamic properties that differ from the average for each protein, we quantified deviations in peak intensities across variants (Figure 2, Supplementary Figure 10). Specifically, we measured differences between each peak intensity and the median intensity for all assigned peaks in the spectrum and normalized these differences by the standard deviation in peak intensities for each variant. According to this calculation, a negative peak intensity deviation results from peaks having intensities that are lower than the average, indicating slower, exchange-dominated dynamics, while positive deviations reflect higher than average intensities due to increased flexibility. In addition, peaks that are similar in intensity to the average are considered rigid relative to these two dynamics regimes. Correlation analysis of intensity deviations across the protein series was then performed for each residue to identify functional dynamics hotspots. 24 residues (Supplementary Table 3) showed a decrease in intensity deviation as the quantum yield of the variant increased (mScarlet < mCherry < mPlum-E16P < mRojoA/mRouge), suggesting that increases in rigidity of these residues gave rise to higher quantum yields. These residues were primarily localized to β-strands 7–10 (Supplementary Figure 11) on the β-barrel face adjacent to the chromophore’s phenolate moiety (termed the “phenolate face”), as well as the N-terminal end of the central α-helix that spans the barrel and contains the chromophore (Figure 3a). Most of these residues are distal to the chromophore, and form a contiguous band across the phenolate face, suggesting a long-range structural network that rigidifies the chromophore through a mechanical pathway. Additionally, a further 24 residues on the opposing “acylimine face” (Supplementary Figure 11) showed an increase in intensity deviations with increasing quantum yields (mRouge/mRojoA < mPlum-E16P < mCherry < mScarlet), suggesting increased flexibility with higher quantum yield (Figure 3a). Although the link between increased flexibility of the acylimine face and higher quantum yield is unclear, we hypothesize that it may result from compensatory entropy effects associated with phenolate-face rigidification. This hypothesis is consistent with observations that distal protein regions often become more dynamic as binding sites or functional regions rigidify ^22^. The differential effects on the phenolate and acylimine faces further suggest that these dynamics are functionally relevant rather than randomly distributed.

**Figure 2.**
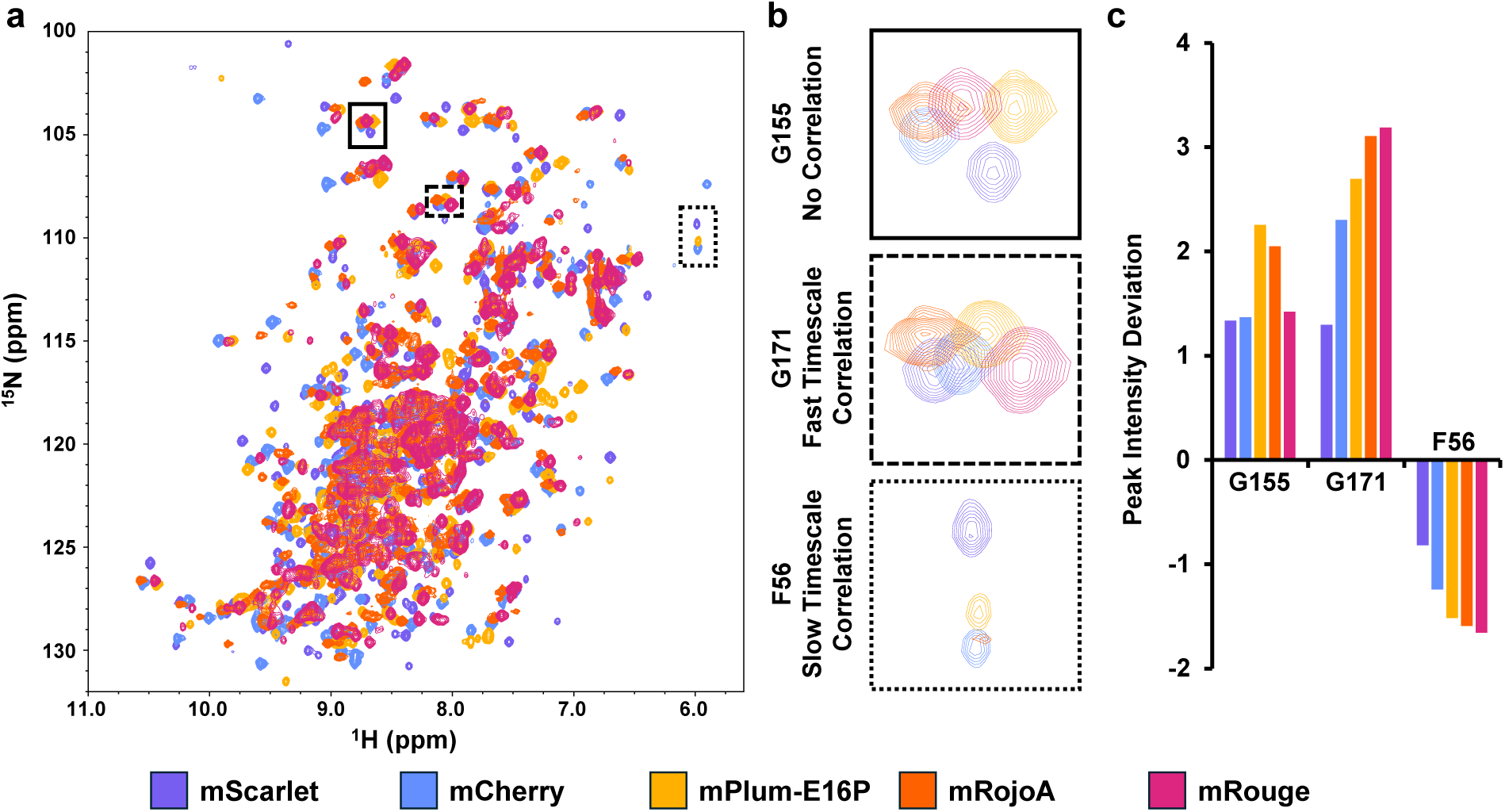
^1^H-^15^N-HSQC peak intensities correlate RFP quantum yield with structural dynamics. (a) Overlay of ^1^H-^15^N-HSQC spectra for five RFPs reveals regions of altered dynamics via differences in peak intensities. Spectra are normalized to a constant median peak intensity for each protein. (b) Insets from (a) showing peaks from residues with no trend in peak intensity differences across RFPs (G155), increased peak intensities in dimmer RFPs (G171) and decreased peak intensities in dimmer RFPs (F56). (c) Deviations in peak intensities for the residues shown in (b). Positive peak intensity deviations result from fast (ps–ns) motions and negative peak intensity deviations reflect slow motions (μs–s). Bars are ordered by descending quantum yield at each residue.

**Figure 3.**
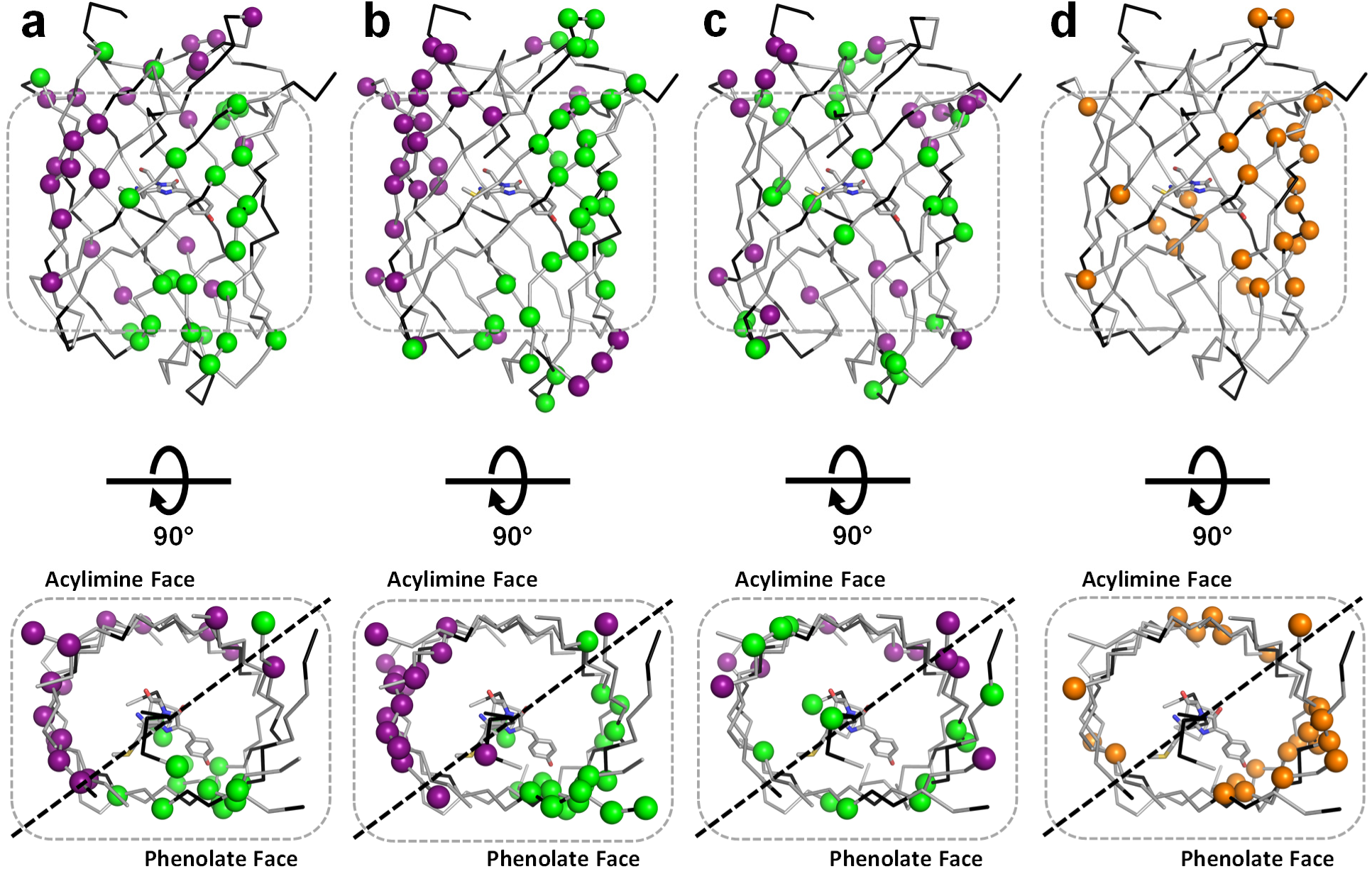
Residue positions linked to quantum yield in RFPs. Green and purple spheres indicate residues whose dynamics are positively or negatively correlated with quantum yield, respectively, based on (a) NMR peak intensity analysis, (b) Z-score analysis of non-cryogenic B-factors, and (c) Z-score analysis of cryogenic B-factors. (d) Orange spheres mark residues mutated in at least two directed evolution campaigns that increased brightness. In all cases, residues are mapped onto the mCherry crystal structure (PDB ID: 2H5Q) ^36^ and the chromophore is shown as sticks.

To assess whether slower conformational exchange contributed to differences in quantum yield, we performed ^15^N CPMG relaxation dispersion experiments to detect motions on the microsecond to millisecond timescale ^23,24^. Neither mCherry nor mRojoA exhibited significant relaxation dispersions (Supplementary Figure 12), suggesting that dynamics on this timescale arising from assignable residues are not substantially functionally relevant, which is consistent with typical RFP fluorescence lifetimes of 1–5 ns (e.g., 1.5 ns for mCherry ^25^, 4 ns for mScarlet ^26^. This observation is further supported by the nature of the HSQC peak intensity correlations we observed. 67% (16 out of 24) of all correlations between rigidity and quantum yield resulted from positive deviations from the mean (Supplementary Figure 10), which are representative of ps–ns timescale dynamics. Moreover, on the phenolate face, this proportion increases to 79% (11 out of 14), hinting that fast motions dominate functional relevance in this region.

To further validate these observations, we measured backbone ^15^N T_1_ and T_2_ relaxation ^27,28^ and calculated apparent correlation times on a per-residue level (Supplementary Figure 13). Deviations from the average were calculated to identify sites undergoing dynamics on this faster timescale, however there were no clear correlations between dynamic residues and quantum yield. This observation partly reflected the difficulty in obtaining adequate signal-to-noise levels across the series of spectra needed for accurate measurements for some of the variants. These results highlight the advantages of NMR peak intensity analysis for identifying regions of the protein that undergo functionally relevant fast-timescale motions, offering greater sensitivity and ease compared to ^15^N spin relaxation measurements. This advantage is especially evident for proteins that are large by solution NMR standards or that exhibit intermediate-timescale dynamics, both of which can broaden peaks and degrade spectral quality due to relaxation during the pulse sequence delays required for relaxation measurements. Although peak intensity analysis provides less detailed characterization of fast backbone dynamics, its higher sensitivity to residues dominated by fast motion appears sufficient to pinpoint functional dynamics hotspots.

### Crystallographic evidence of dynamics hotspots

To validate the functional dynamics hotspots identified by peak intensity analysis, we turned to non-cryogenic X-ray crystallography as an orthogonal approach. Although there is always the potential for crystal packing to limit conformational flexibility ^29–31^, it is generally appreciated that non-cryogenic crystal structures provide a more accurate representation of a protein’s conformational ensemble compared to those determined using cryogenic data ^32–34^. It has also been demonstrated that B-factors and alternative side-chain conformations modeled from crystallographic data collected at non-cryogenic temperatures contain information about protein dynamics that correlates with NMR-derived order parameters that probe fast (ps–ns) motions ^35^.

We solved crystal structures of mCherry and two variants—a dim variant (mRouge) and a bright variant (mScarlet) (Supplementary Table 4)—at 277 K. Although mCherry and mRouge crystallized in the P 1 21 1 space group and mScarlet in C 1 2 1, all structures contained a single molecule per asymmetric unit (Supplementary Table 5). To evaluate whether the correlations between rigidity and quantum yield observed by NMR could be observed in the crystal structures, we compared per-residue B-factors across structures, using Z-score normalization to account for differences between crystal forms (Supplementary Figure 14–16). Consistent with NMR results (Figure 3a), residues on the phenolate face exhibited reduced mobility with increasing quantum yield, while those on the acylimine face became more flexible (Figure 3b). These findings support the hypothesis that mutations enhancing quantum yield act by rigidifying the phenolate face while allowing increased flexibility on the opposing side, and in turn further validate the use of HSQC peak intensities to report broadly on protein dynamics in fluorescent proteins.

Notably, the trends relating rigidity of the phenolate face and flexibility of the acylamine face with increased quantum yield were absent when the same analysis was performed on cryogenic structures previously deposited into the PDB (Figure 3c), despite the fact that the cryogenic (PDB ID: 2H5Q ^36^, 3NED ^20^, 5LK4 ^19^) and non-cryogenic structures share the same space group and have similar unit cell dimensions. Specifically, when quantifying overlap between datasets (Supplementary Figure 17), we observe roughly twice as many exact matches between our NMR results and non-cryogenic crystallography than with cryogenic crystallography. Additionally, analysis of non-cryogenic crystal structures and NMR intensities show spatial clustering of residues whose dynamics correlate with quantum yield (Figure 3). In contrast, analysis of cryogenic crystal structures reveals a much more dispersed pattern of residues whose dynamics correlate with quantum yield. While there is generally good agreement between analyses performed on non-cryogenic crystal structures and on NMR intensities, >50% of the correlations found by analyzing cryogenic crystal structures cannot be matched to either the NMR or non-cryogenic crystallography datasets, further highlighting the lack of clustering observed in this dataset.

### Functional relevance of the identified hotspots

Having identified and validated dynamics hotspots, we next assessed their functional relevance by analyzing mutation patterns from multiple directed evolution campaigns aimed at enhancing RFP brightness ^16,19,20,37–46^, which is directly proportional to quantum yield. These campaigns produced beneficial mutations distributed across the protein scaffold, both in direct contact with the chromophore and at distant sites. Given the known relationship between chromophore dynamics and quantum yield, we hypothesized that brightness-enhancing mutations would localize to regions identified in our NMR analysis—namely, the phenolate face and the central α-helix.

We analyzed 14 independently developed RFPs that underwent brightness optimization by protein engineering (Supplementary Table 6). To minimize noise from random mutagenesis, we focused on positions mutated in at least two separate studies and mapped them onto the RFP structure (Figure 3d). Notably, 55% of these recurrent mutations also localized to β-strands 7–10, reinforcing the importance of the phenolate face in controlling chromophore brightness. However, while mutations cluster primarily at the centers of β-strands 8 and 9, our NMR results highlight the termini of strands 8–10 and the center of strand 7 as key sites for functionally relevant dynamics. Although these sites are proximal to mutated clusters, the overlap is not exact. This is not unexpected, as NMR chemical shifts report on aggregate motions that give rise to differences in the electronic environment of a nucleus rather than on motion of the specific nucleus itself. Similarly, mutations are liable to affect dynamical motions not only at the backbone of the mutated residue but in nearby interacting positions as well. We note here that the central residues of strand 8 could not be assigned in the lower-quality mRouge and mRojoA spectra and were therefore excluded from our analysis. However, the inability to assign a residue is usually a consequence of substantial line broadening, suggesting that these residues are also involved in significant conformational exchange in the μs to ms timescale. This is further corroborated by our crystallography data, where additional positions in this unassigned region at the center of strand 8 were highlighted as key dynamic sites.

It is important to note that perfect overlap between functional dynamics hotspots and the mutational database should not be expected since mutant selection is also influenced by other factors that contribute to brightness, such as chromophore maturation rates. Chromophore maturation requires two oxidation steps ^47^ and is partially dependent on oxygen access through a channel formed between the ends of β-strands 7 and 10 ^48^. Mutations that overly rigidify this region may enhance quantum yield but hinder oxygen access, delaying chromophore maturation. As a result, mutations that enhance quantum yield but delay maturation are less likely to be retained during high-throughput screening of RFP libraries for enhanced brightness.

The functional relevance of the dynamics hotspots we identified is underscored by prior findings from our engineering of mSandy2 ^16^, a bright RFP derived from mRojoA. Guided by the principle that increased chromophore rigidity enhances quantum yield, we used computational design with the Triad software ^20,49,50^ to tighten packing around the chromophore phenolate. This rational design process introduced three mutations on the phenolate face and one on the central helix, yielding mSandy1, a variant with a 13-fold increase in quantum yield (0.26). Subsequent directed evolution using random mutagenesis produced mSandy2 (quantum yield 0.36), which incorporated five additional phenolate-face mutations (Supplementary Figure 18). Three of these mutations were distal to the chromophore, outside the first shell of direct contact supporting the conclusion that brightness can be modulated by dynamic regions beyond the immediate chromophore environment.

## Discussion

Our results establish NMR peak intensity analysis as a useful and accessible tool for identifying functionally relevant dynamics hotspots. NMR peak lineshape analyses have historically provided insight into dynamics on the 10 to 100 millisecond timescale ^10,12,14^. However, to obtain dynamics information beyond this restricted kinetic window, it is necessary to perform specific NMR experiments that are tailored to a target timescale (e.g. CPMG relaxation dispersion for µs–ms timescale dynamics and nuclear spin relaxation measurements for ps–ns dynamics). Acquisition and analysis of these datasets require substantially more instrument and analysis time relative to that needed for the acquisition and analysis of the single ^15^N-HSQC spectrum used in peak intensity analysis. Moreover, the lower inherent sensitivity of these dynamics experiments makes it difficult to acquire complete datasets for larger proteins (>30 kDa), and/or proteins with spectra exhibiting μs–ms timescale peak broadening. In fact, sensitivity losses complicated comparison of dynamics across the RFP series, partly due to the molecular weights being near 30 kDa and the substantial peak broadening observed in spectra from dimmer variants. Although ^15^N spin relaxation experiments could be successfully acquired for all variants, there were several residues that could not be analyzed in these experiments, limiting the number of correlations that could be made across the RFP series. While our peak intensity analysis method did not provide a quantitative measurement of exchange rates, the higher degree of sequence coverage and ability to distinguish sites dominated by fast (ps–ns) versus slow (ms) timescale dynamics provides more useful insight into the localized dynamics that may be important for function, as we observed for the RFPs.

We anticipate that the peak intensity analysis approach presented here will be broadly applicable to any protein family for which assignable spectra can be obtained. This method can also be extended to proteins larger than 30 kDa by employing deuteration at non-exchangeable sites and using relaxation-optimized pulse sequences on high-field NMR spectrometers ^51,52^. The simplicity of peak intensity analysis makes it particularly attractive for protein engineering workflows, as a single assigned spectrum of the wild-type protein is sufficient to identify dynamic regions suitable for saturation mutagenesis and downstream screening. Subsequent analyses of variants with enhanced or reduced function can then guide successive rounds of dynamics-informed engineering. Notably, this approach does not require prior knowledge of the timescales of functionally relevant motions, in contrast to conventional NMR relaxation methods that probe specific exchange regimes.

This strategy complements other applications of NMR in protein engineering. For example, Korendovych and colleagues ^4^ used NMR chemical shift perturbations (CSPs) to identify regions affected by ligand binding in a *de novo* enzyme scaffold. Saturation mutagenesis of residues outside the active site that exhibited significant CSPs yielded variants with enhanced catalytic efficiency. Historically, CSP analyses have been used to pinpoint substrate- or ligand-binding hotspots ^5,53^. Because these regions are often strongly coupled to function, CSPs provide an accessible means to identify first-shell residues for engineering new or improved enzyme activities. However, fine-tuning protein properties frequently requires mutations beyond these first-shell residues ^54,55^, where more subtle dynamical and epistatic effects come into play ^56^, factors that are often challenging to predict or measure experimentally. The NMR peak intensity analysis described here enables the identification of regions undergoing fast or slow local motions as potential sites of functional importance outside the immediate binding site. As with CSPs, this approach does not require high-resolution structural information, allowing rapid assessment of new protein systems and their dynamics.

In summary, our findings establish a framework for integrating readily accessible dynamics data into protein engineering workflows. Acquiring HSQC spectra across a panel of variants with different activities allows efficient transfer of resonance assignments once a single spectrum has been assigned, a process that can be further streamlined with deep learning-based tools ^57^. When variants are unavailable, random mutagenesis can generate suitable candidates, and even low-activity variants yield valuable comparative information. Peak intensity analysis can then pinpoint residues where local rigidity or flexibility correlates with function, guiding saturation mutagenesis toward activity-enhancing substitutions. Beneficial mutations can be combined to achieve additive or synergistic improvements, while ongoing NMR monitoring helps refine the search space and direct iterative optimization.

Because this approach requires only backbone amide assignments from a small number of variants, it offers a straightforward and generalizable route for integrating structural dynamics into protein engineering campaigns. Importantly, the method is experimentally simple, broadly applicable to soluble proteins up to ∼300 residues, and capable of probing a wide range of motion timescales, including the millisecond dynamics often critical for enzyme catalysis ^9,58^. We therefore anticipate that this NMR-guided workflow will extend beyond fluorescent proteins to diverse systems, enabling the rational tuning of dynamics to enhance a broad spectrum of protein functions.

## Methods

### Red fluorescent protein genes

Amino-acid sequences for all RFPs described here are listed on Supplementary Table 1. His-tagged (N-terminus) genes for mCherry, mRouge, mRojoA and mPlum-E16P cloned into pET-11a (Novagen) were a gift from Stephen L. Mayo ^20,21^. The mScarlet gene was obtained from Addgene (pCytERM_mScarlet_N1; a gift from Dorus Gadella – Addgene plasmid #85066; http://n2t.net/addgene:85066; RRID: Addgene_85066) and cloned into the pET-11a vector via *Nde*I and *Bam*HI. All plasmids were transformed into *E. coli* BL21-Gold (DE3) cells for protein expression. All open-reading frames were verified by DNA sequencing.

### Protein expression and purification

RFP variants were expressed in *E. coli* BL21-Gold (DE3) cells (Agilent) using lysogeny broth (LB) supplemented with 100 µg mL^−1^ ampicillin. Cells were grown at 37°C with shaking until they reached an optical density at 600 nm of 0.6. Protein expression was induced by adding 1 mM isopropyl β-D-1-thiogalactopyranoside (Thermo Scientific) and cultures were incubated overnight at 16 °C with shaking. Cells were then pelleted by centrifugation and resuspended in 8 mL lysis buffer (100 mM potassium phosphate buffer, pH 7.4, 5 mM imidazole, 1 mg mL^−1^ lysozyme, 50 U benzonase nuclease [Novagen]). Cells were lysed using an EmulsiFlex-B15 cell disruptor (Avestin) and lysates were harvested by centrifugation. Proteins were purified by Ni-NTA affinity chromatography using Econo-Pac chromatography columns (Bio-Rad) according to the manufacturer’s protocol. Eluted fractions were desalted using Macrosep Advance centrifugal devices (Pall) into phosphate-buffered saline (137 mM sodium chloride, 2.7 mM potassium chloride, 10 mM disodium phosphate, 1.8 mM monopotassium phosphate, pH 7.4). Protein samples were stored at 4°C for two days to allow for chromophore maturation prior to spectroscopic characterization. For crystallography experiments, an additional purification step consisting of gel filtration into 20 mM sodium phosphate buffer pH 7.4 was performed using an ÄKTA pure (GE Healthcare) fast protein liquid chromatography system equipped with a Superdex 75 (GE Healthcare) column. Protein purity was verified by SDS-PAGE.

For NMR experiments, RFP variants were expressed in M9 minimal medium supplemented with 1 g L^‒1^ ^15^N-ammonium chloride and/or 3 g L^‒1^ ^13^C-D-glucose for isotopic labeling. Cultures were grown at 37 °C with shaking to an OD₆₀₀ of approximately 0.6, at which point expression was induced with 1 mM isopropyl-β-D-thiogalactopyranoside. After overnight incubation at 16 °C (for mRojoA, mRouge, and mPlum-E16P) or 37 °C (for mCherry and mScarlet), cells were harvested by centrifugation and lysed using an EmulsiFlex-B15 cell disruptor (Avestin). Proteins were purified by immobilized metal affinity chromatography (Qiagen) followed by size-exclusion chromatography on an ENrich SEC 650 column (Bio-Rad) in 10 mM sodium phosphate buffer (pH 7.4). Final samples were concentrated using Amicon Ultracel-3K (EMD Millipore) or Macrosep Advance 10K (Pall) centrifugal filter units.

### Spectroscopic characterization

Absorption, excitation (λ_em_ = 655 nm) and emission (λ_ex_ = 535 nm) spectra were measured in phosphate-buffered saline (137 mM sodium chloride, 2.7 mM potassium chloride, 10 mM disodium phosphate, 1.8 mM monopotassium phosphate, pH 7.4) using an Infinite M1000 (Tecan) plate reader. Quantum yields were extrapolated by comparing the integrated fluorescence intensity of the mutant proteins with that of equally absorbing samples of mCherry and mRaspberry (quantum yields of 0.23 and 0.15, respectively) with excitation at 535 nm. Extinction coefficients were determined using the method described by Kredel and colleagues ^59^, which takes into account the presence of green chromophore in the population of RFP molecules, giving a more accurate estimate. Briefly, protein samples were diluted (1:10) into Britton-Robinson buffers ^60^ (pH 11–13) to allow measurable slow denaturation of the native red chromophore (peak at 568–605 nm, depending on the RFP) to the green form (452 nm). From the absorption spectrum recorded at different time intervals, a ratio of extinction coefficients of the native chromophore species to that of the denatured species (44,000 M^−1^ cm^−1^) was calculated, from which the value for the red chromophore was extrapolated.

### NMR spectroscopy

^15^N- and ^13^C-labeled RFP samples for NMR were prepared at 0.1–0.2 mM in 10 mM sodium phosphate buffer (pH 7.4), supplemented with 10 µM EDTA, 0.02% sodium azide, 1× cOmplete EDTA-free Protease Inhibitor Cocktail (Roche), and 5% D_2_O. All NMR experiments were performed at 25 °C on a Bruker AVANCEIII HD 600 MHz spectrometer equipped with a triple resonance cryoprobe. Data were processed using NMRPipe and analyzed with NMRViewJ ^61^. Backbone chemical shift assignments were obtained using standard 3D triple resonance experiments: HSQC, HNCO, HNCACB, and CBCA(CO)NH ^62^. HSQC spectra used for peak intensity analyses were measured using a phase-sensitive pulse sequence using water flip-back and WATERGATE selective pulse ^63–66^ with 64 scans, an interscan delay of 5 seconds, and 256 points in the indirectly detected dimension. ^15^N T_1_ and T_2_ relaxation times were measured using gradient enhanced sensitivity-based HSQC experiments ^67^ with 12 time points spanning 10–2000 ms, 98 scans, an interscan delay of 1 second, and 256 points in the indirectly detected dimension for T_1_ measurements and 16.96 to 271.36 ms, 104 scans, an interscan delay of 1 second, and 256 points in the indirectly detected dimension for T_2_ measurements, each run in a randomly shuffled order. ^15^N CPMG relaxation-dispersion spectra were acquired in a constant time manner^68^ using eleven points with frequencies ranging from 10 to 1000 Hz, run in a randomly shuffled order.

### Protein crystallization

Purified RFPs were concentrated using Amicon Ultra-15 3K centrifugal devices (Millipore Sigma) to a concentration of 10–20 mg mL^−1^ in 200 mM sodium phosphate buffer pH 7.4. Crystallization drops were prepared by mixing 1 µL of protein solution with 1 µL of the mother liquor and sealed inside a reservoir containing an additional 500 µL of the mother liquor solution. Crystallization conditions used to produce the samples used for used for X-ray data collection are provided in Supplementary Table 4.

### X-ray data collection and processing

Prior to X-ray data collection, crystals were mounted on polymer MicroMounts (MiTeGen) and sealed using a MicroRT tubing kit (MiTeGen). Single-crystal X-ray diffraction data was collected on beamline 8.3.1 at the Advanced Light Source. The beamline was equipped with a Pilatus3 S 6 M detector and was operated at a photon energy of 11111 eV. Crystals were maintained at 277 K throughout the course of data collection. Each data collection scan included 360 images covering a 180° wedge of reciprocal space. Scans were collected using a minimal X-ray dose to avoid radiation damage to the chromophores. For mRouge and mScarlet, each data set was collected using an X-ray dose of 22.3 kGy, and for mCherry, each data set was collected using an X-ray dose of 17.3 kGy. Multiple scans were collected for each RFP variant either from different crystals, or if their size permitted, from unique regions of single large crystals.

X-ray data were processed with the Xia2 program ^69^, which performed indexing and integration with DIALS ^70^, followed by scaling and merging with DIALS.SCALE ^71^. For mRouge and mScarlet, individual scans were sufficient to yield high-quality data sets. For mCherry, we merged eight individual scans to generate a single, reduced data set.

### Structure determination

We obtained initial phase information for calculation of electron density maps by molecular replacement using the program Phaser ^72^, as implemented in the PHENIX suite ^73^, with the crystal structure of mCherry (PDB ID: 2H5Q) used as search model. We rebuilt the initial models using the electron density maps calculated from molecular replacement. We then performed iterative refinement of atomic positions, individual atomic displacement parameters (B-factors), and occupancies using a riding-hydrogen model and TLS-based automatic weight optimization until convergence was achieved, except for mCherry, where TLS was not applied. All model building was performed using Coot 0.8.9.256 ^74^ and refinement steps were performed with phenix.refine (v1.21.5207) within the PHENIX suite ^73,75^. Restraints for the chromophore were generated using phenix.elbow ^76^, starting from coordinates available in the Protein Data Bank (PDB ligand ID: NRQ). Further information regarding model building and refinement is presented in Supplementary Table 5.

### Dynamics analysis

HSQC peak intensities were deconvoluted and fit to Gaussian models using NMRDraw ^77^. Peak intensities were converted to a deviation metric by calculating the difference between each peak intensity and the median intensity for that variant, then normalizing these differences by the standard deviation for each variant. Residues were classified as having rigidity positively correlated with quantum yield if the absolute intensity deviation followed the order: mScarlet < mCherry < mPlum-E16P < mRouge or mRojoA. Rigidity negatively correlated with quantum yield followed the reverse order. mRojoA and mRouge were not directly compared due to similar quantum yields. Additionally, a threshold of ≥ 0.5 standard deviations for mRouge or mRojoA for positive correlations, or mScarlet for negative correlations was applied to avoid false positives from uniformly rigid positions.

*T_1_-T_2_* relaxation experiments were analyzed using the Bruker Dynamics Center software, by fitting a Gaussian model to peaks and fitting peak intensities over 12-point relaxation curves using non-linear regression. Correlation time measurements were derived from *T_1_* relaxation and *T_2_* relaxation values, calculated using the approximation 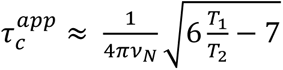 and converted to a normalized correlation time metric by dividing correlation times for each residue by the median correlation time for that protein to remove error introduced by variation in the global protein tumbling rate between proteins.

In the analysis of X-ray crystallographic models, refined B-factors were Z-score normalized using the following equation to facilitate comparison across different crystal structures: 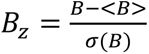. Residues were classified as having rigidity positively correlated with brightness if the B-factor Z-score followed the order: mScarlet < mCherry < mRouge. Rigidity negatively correlated with brightness followed the reverse order. Additionally, to avoid false positives from statistical noise, residues with a B-factor Z-score difference between mScarlet and mRouge of < 0.3 were discarded, which corresponds roughly to 0.5 standard deviations for these datasets.

## Acknowledgements

R.A.C. acknowledges grants from the Canada Foundation for Innovation (26503), Natural Sciences and Engineering Research Council of Canada (RGPIN-2016-04831), and the Human Frontier Science Program (RGP0041/2016). A.M.D. is the recipient of an Ontario Graduate Scholarship and a postgraduate scholarship from the Natural Sciences and Engineering Research Council of Canada (NSERC). S.E.H. is the recipient of an undergraduate student research award from NSERC. S.L. is the recipient of an Ontario Graduate Scholarship and a postgraduate scholarship from NSERC. We thank Stephen L. Mayo for providing the mCherry, mPlum-E16P, mRouge and mRojoA expression vectors.

## Author Contributions

A.M.D. and R.A.C. conceived the project. A.M.D. and S.E.H. performed NMR experiments. A.M.D. and S.E.H. analyzed NMR data. N.K.G. designed NMR experiments. S.L. performed spectral characterization of RFPs. A.M.D., S.E.H., S.L. purified proteins. M.C.T. crystallized proteins and performed X-ray diffraction experiments. R.A.C. performed model building and refinements. A.M.D. and R.A.C. wrote the manuscript. M.C.T. and N.K.G. edited the manuscript.

## Data Availability

Chemical shift assignments have been deposited in the Biological Magnetic Resonance Data Bank with accession codes 27906 (mCherry), 27907 (mPlum-E16P), 27908 (mScarlet), 27909 (mRouge), and 27910 (mRojoA). Structure coordinates for red fluorescent proteins have been deposited in the RCSB Protein Data Bank with the following accession codes: mRouge (PDB ID: 9YVX), mCherry (PDB ID: 9YVY), and mScarlet (PDB ID: 9YVZ).

