## Supplementary Information for "Mapping Functional Dynamics Hotspots for Protein Engineering with NMR Peak Intensity Analysis"

**This file includes:**

Supplementary Tables 1–6

Supplementary Figures 1–16

**Supplementary Table 1.** Amino-acid sequences of various RFPs

| Protein | Molecular weight (kDa) | Sequence [a] |
|---|---|---|
| **mScarlet** | 27.2 | MHHHHHHGVSKGEAVIKEFMRFKVHMEGSMNGHEFEIEGEGEGRPYEGTQTAKLKVTKGGPLPFSWDIL<br>SPQFMYGSRAFTKHPADIPDYYKQSFPEGFKWERVMNFEDGGAVTVTQDTSLEDGTLIYKVKLRGTNFP<br>PDGPVMQKKTMGWEASTERLYPEDGVLKGDIKMALRLKDGGRYLADFKTTYKAKKPVQMPGAYNVDRKL<br>DITSHNEDYTVVEQYERSEGRHSTGGMDELYK |
| **mCherry** | 27.7 | MGHHHHHHGVSKGEEDNMAIIKEFMRFKVHMEGSVNGHEFEIEGEGEGRPYEGTQTAKLKVTKGGPLPF<br>AWDILSPQFMYGSKAYVKHPADIPDYLKLSFPEGFKWERVMNFEDGGVVTVTQDSSLQDGEFIYKVKLR<br>GTNFPSDGPVMQKKTMGWEASSERMYPEDGALKGEIKQRLKLKDGGHYDAEVKTTYKAKKPVQLPGAYN<br>VNIKLDITSHNEDYTIVEQYERAEGRHSTGGMDELYK |
| **mPlum-E16P** | 26.5 | MGHHHHHHGVSKGEEVIKEFMRFKPHMEGSVNGHEFEIEGEGEGRPYEGTQTARLKVTKGGPLPFAWDI<br>LSPQIMYGSKAYVKHPADIPDYLKLSFPEGFKWERVMNFEDGGVVTVTQDSSLQDGEFIYKVKVRGTNF<br>PSDGPVMQKKTMGWEASSERMYPEDGALKGEMKMRLRLKDGGHYDAEVKTTYMAKKPVQLPGAYKTDIK<br>LDITSHNEDYTIVEQYERAEGRHSTGA |
| **mRojoA** | 27.7 | MGHHHHHHGVSKGEEDNMAIIKEFMRFKTHMEGSVNGHEFEIEGEGEGRPYEGTQTAKLKVTKGGPLPF<br>AWDILSPQFMYGSKAYVKHPADIPDYLKLSFPEGFKWERVMNFEDGGVVTVTQDSSLQDGEFIYKVKLH<br>GTNFPSDGPVMQKKTMGWEASSERMYPEDGALKGEIKLRLKLKDGGHYDAEVKTTYKAKKPVQLPGAYN<br>ANYKLDITSHNEDYTIVEQYERCEGRHSTGGMDELYK |
| **mRouge** | 27.7 | MGHHHHHHGVSKGEEDNMAIIKEFMRFKTHMEGSVNGHEFEIEGEGEGRPYEGTQTAKLKVTKGGPLPF<br>AWDILSPQFMYGSKAYVKHPADIPDYLKLSFPEGFKWERVMNFEDGGVVTVTQDSSLQDGEFIYKVKLR<br>GTNFPSDGPVMQKKTMGWEACSERMYPEDGALKGEMKMRLKLKDGGHYDAEVKTTYKAKKPVQLPGAYN<br>TNTKLDITSHNEDYTIVEQYERNEGRHSTGGMDELYK |

[a] All RFPs contain a His-tag at the N-terminus.

**Supplementary Table 2.** Mutations in RFP variants

| **Protein** | **# of mutations from mCherry [a]** | **Mutations from mCherry [b]** |
|---|---|---|
| mScarlet | 30 | I7V V22M A57S K70R Y72F V73T L83Y L84Q V104A S111T Q114E E117T F118L S131P S147T M150L A155V E160D Q163M R164A K166R H172R D174L E176D V177F L189M N196D I197R I210V A217S |
| mPlum-E16P | 13 | A6E I7V V16P K45R F65I L124V I161M Q163M K166R K182M N194K V195T N196D |
| mRojoA | 6 | V16T Q163L R125H V195A I197Y A217C |
| mRouge | 7 | V16T S146C I161M Q163M V195T I197T A217N |

[a] Does not include N- and C-terminal tags nor associated linker sequences.
[b] Numbering based on the mCherry crystal structure (PDB ID: 2H5Q).

**Supplementary Table 3.** RFP residues with dynamics correlated to quantum yield

| Method | Positive correlation [a] | Negative correlation [b] |
|---|---|---|
| NMR peak intensity analysis | 8, 24, 56, 57, 59, 60, 84, 116, 128, 140, 142, 144, 145, 146, 156, 165, 167, 168, 171, 196, 197, 214, 216, 219 | 15, 19, 21, 32, 33, 34, 35, 43, 45, 48, 78, 81, 82, 86, 90, 104, 106, 113, 114, 158, 182, 187, 188, 210 |
| B-factor Z-scores, 277 K | 56, 57, 59, 76, 77, 78, 79, 132, 133, 139, 141, 143, 144, 145, 146, 147, 161, 174, 176, 177, 180, 188, 195, 196, 197, 206, 217, 218, 219, 220, 221 | 8, 9, 10, 11, 12, 13, 14, 15, 16, 23, 30, 32, 34, 36, 48, 49, 72, 114, 115, 118, 120, 121, 155, 156, 169, 170, 171, 187, 207, 210 |
| B-factor Z-scores, 100 K | 44, 51, 52, 65, 71, 85, 102, 112, 113, 121, 130, 131, 133, 134, 146, 151, 161, 162, 168, 184, 185, 186, 196, 200, 206, 216 | 9, 10, 11, 21, 29, 49, 104, 105, 116, 117, 129, 154, 156, 157, 158, 171, 187, 192, 193, 207, 208 |

[a] Residue positions where quantum yield increases with increasing rigidity. Numbering based on the mCherry crystal structure (PDB ID: 2H5Q).
[b] Residue positions where quantum yield increases with increasing flexibility. Numbering based on the mCherry crystal structure (PDB ID: 2H5Q).

**Supplementary Table 4.** Crystallization conditions

| **Protein** [a] | **Concentration ($mg\ mL^{-1}$)** | **Mother liquor** | **Soak** [b] |
|---|---|---|---|
| **mRouge** | 15 | 0.1 M Tris, pH 8.3<br>0.25 M $MgCl_2$<br>20% PEG-4K | 0.1 M HEPES, pH 7.5<br>0.25 M $MgCl_2$<br>20% PEG-4K |
| **mCherry** | 10 | 0.1 M Tris, pH 8.5<br>0.1 M sodium acetate<br>30% PEG-4K | – |
| **mScarlet** | 20 | 0.15 M $(NH_4)_2SO_4$<br>24% PEG-3350 | 0.1 M HEPES, pH 7.5<br>0.15 M $(NH_4)_2SO_4$<br>24% PEG-3350 |

[a] All proteins were crystallized at 20 °C.
[b] Soaking was done to change pH of crystals to 7.5.

**Supplementary Table 5.** Crystallographic data and refinement statistics

| **Name** | **mRouge** | **mCherry** | **mScarlet** |
|---|---|---|---|
| PDB ID | 9YVX | 9YVY | 9YVZ |
| **Data collection** [a] | | | |
| Temperature (K) | 277 | 277 | 277 |
| X-ray dose (kGy) | 22.3 | 17.3 | 22.3 |
| Resolution (Å) | 45.65–1.38 | 57.94–1.65 | 58.38–1.35 |
| Space group | P 1 21 1 | P 1 21 1 | C 1 2 1 |
| *Cell params.* | | | |
| a b c (Å) | 49.19 43.49 61.78 | 48.94 43.86 61.73 | 84.13 48.54 59.86 |
| α β γ (°) | 90.0 111.9 90.0 | 90.0 110.7 90.0 | 90.0 102.8 90.0 |
| $R_{pim}$ | 0.063 (0.379) | 0.065 (0.433) | 0.071 (0.325) |
| $CC_{1/2}$ | 0.990 (0.668) | 0.995 (0.643) | 0.985 (0.714) |
| I/σI | 6.1 (1.5) | 6.1 (0.7) | 5.5 (1.6) |
| Completeness (%) | 96.7 (91.0) | 98.9 (78.2) | 98.7 (83.7) |
| Multiplicity | 3.3 (3.1) | 19.4 (19.2) | 3.2 (2.5) |
| Wilson B-factor ($Å^2$) | 10.781 | 14.340 | 9.990 |
| # unique reflections | 48458 (2292) | 29217 (1165) | 51150 (2130) |
| **Refinement** | | | |
| R work/free | 0.1253/ 0.1512 | 0.1635/ 0.1828 | 0.1212/ 0.1462 |
| Chains per asymm. unit | 1 | 1 | 1 |
| *No. atoms* | | | |
| Protein | 2031 | 1884 | 2211 |
| Ligand (including chromophore) | 25 | 24 | 33 |
| Water | 209 | 143 | 194 |
| *Averaged B-factors ($Å^2$)* | | | |
| Protein | 18.18 | 21.31 | 17.80 |
| Ligands | 17.34 | 18.02 | 21.99 |
| Water | 31.97 | 29.46 | 30.04 |
| *RMSD* | | | |
| bond lengths (Å) | 0.009 | 0.007 | 0.006 |
| bond angles (°) | 0.840 | 1.089 | 0.890 |
| *Molprobity statistics* | | | |
| Ramachand. outliers (%) | 0.00 | 0.00 | 0.00 |
| Ramachand. allowed (%) | 0.93 | 1.38 | 1.39 |
| Ramachan. favored (%) | 99.07 | 98.62 | 98.61 |
| Rotamer outliers (%) | 0.00 | 0.00 | 0.00 |
| MolProbity clashscore | 3.19 | 2.64 | 3.56 |

[a] Highest resolution shell is shown in parentheses.

**Supplementary Table 6.** Residues mutated during evolutionary campaigns to enhance RFP brightness

| RFP | Brightness [a] ($M^{-1}$ $cm^{-1}$) | Parent | Parent Brightness ($M^{-1}$ $cm^{-1}$) | Mutations [b] | Reference |
|---|---|---|---|---|---|
| **mScarlet** | 70,000 | mCherry [c] | 20,000 | I8V V23M A58S K71R Y73F V74T L84Y L86Q V105A S112T Q115E E118T F119L S132P S148T M151L A157V E161D Q164M R165A K167R H173R D175L E177D V178F L190M N197D I198R I211V A218S | 19 |
| **mScarlet3** | 78,000 | mScarlet-2A-84W | 68,000 | K48R T109A K183R Y194F | 46 |
| **mCherry-XL** | 50,000 | mCherry | 20,000 | W143S I161V Q163Y I197R | 45 |
| **mRojo-VHSV** | 4,700 | mRojoA | 1,600 | T16V P63H W143S L163V | 41 |
| **mRojo-VSHVLF** | 5,400 | mRojo-VHSV | 4,700 | M150L L165F | This study. |
| **mCherry-AYC** | 5,000 | mCherry-I197Y | 2,300 | V195A A217C | 20 |
| **mKate2** | 25,000 | mKate | 15,000 | V45A M146T S158A K231R | 37 |
| **mNeptune2** | 21,000 | mNeptune | 13,000 | A104V I121L I171H K207N | 38 |
| **mRuby3** | 58,000 | mRuby2 | 43,000 | N33R M36E T38V K74A G75D M105T C114E H118N Q120K H159D M160I S171H S173N I192V L202I M209T F210Y H216V F221Y A222S G223N | 39 |
| **mMaroon1** | 8,800 | Maroon0.1 | 5,500 | M11T M15L E16T H23Y K25E S28A G41N E47R T73P Q74P G75D F79Y V93T T95V V101T V104A K120Q L121V V124E G153S C158L N173R K175E V195I | 40 |
| **mCarmine** | 5,800 | mNeptune684 | 1,200 | C61S T103K A104V T105K H157Y P159T I171Q C172T N173F | 42 |
| **mKelly2** | 7,700 | mKelly1 | 7,000 | H72Y T146Y V155E R157T K192E Y193H | 43 |
| **FusionRed-MQV** | 76,000 | FusionRed-M | 24,000 | M41Q C158V | 44 |
| **mSandy2** | 28,000 | mSandy1 | 21,000 | R125H L163V L165F D174V N194Y L199M | 16 |

[a] Brightness is defined as the product of the quantum yield and extinction coefficient.
[b] Mutations are indicated relative to the parent and numbering used in the seminal reference for the engineered RFP.
[c] Although mScarlet was evolved from the non-fluorescent synthetic protein mRed7, mRed7 is itself derived from mCherry.

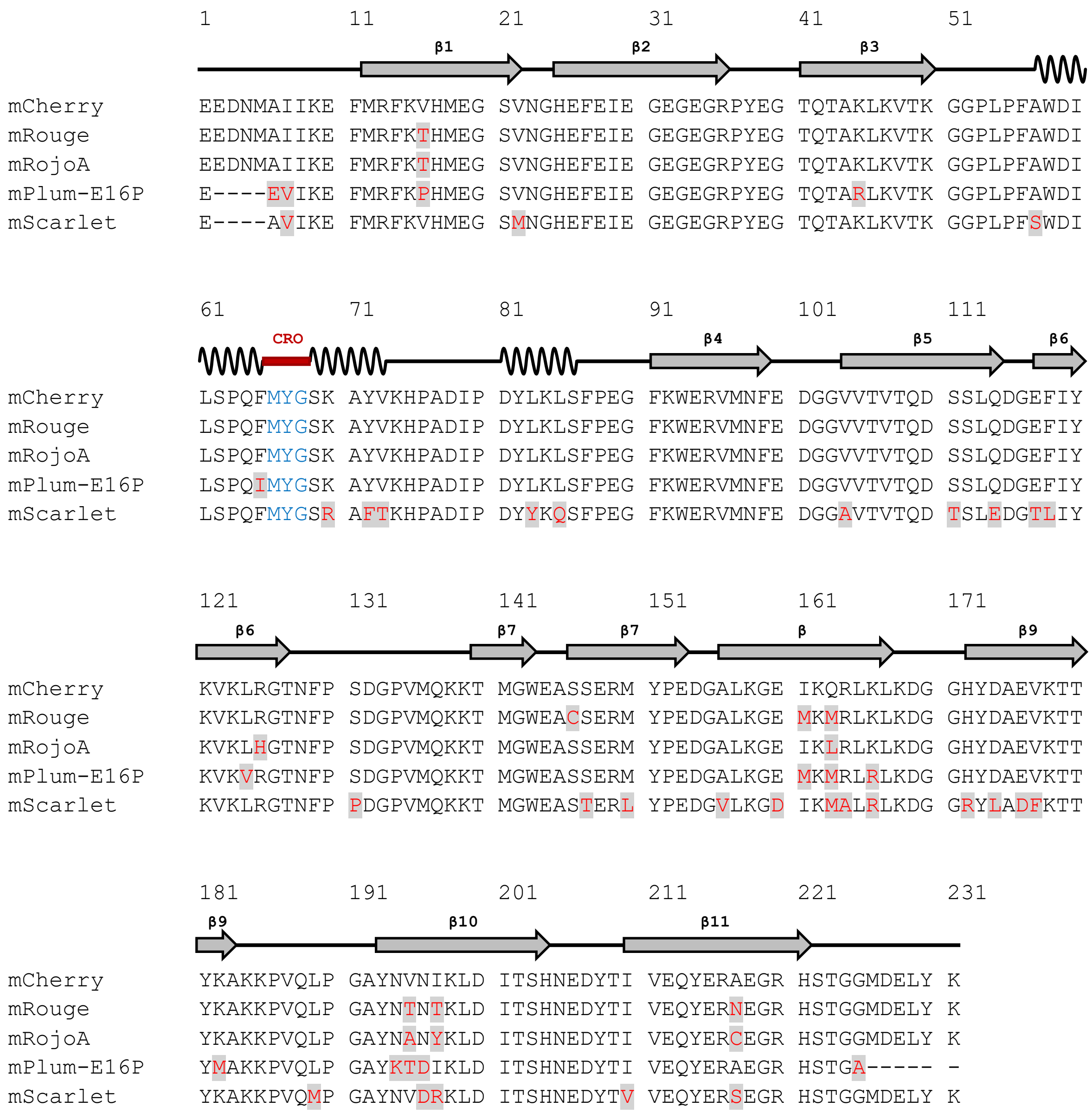


**Supplementary Figure 1. Sequence alignment of RFPs derived from mCherry.** Sequences for mCherry derivatives are shown aligned to the mCherry sequence. Amino acid numbering relative to mCherry. Point mutations are shown in red with grey shading, and the chromophore-forming amino acids (CRO) are shown in blue. mCherry secondary structure elements are also shown aligned to the primary sequences.

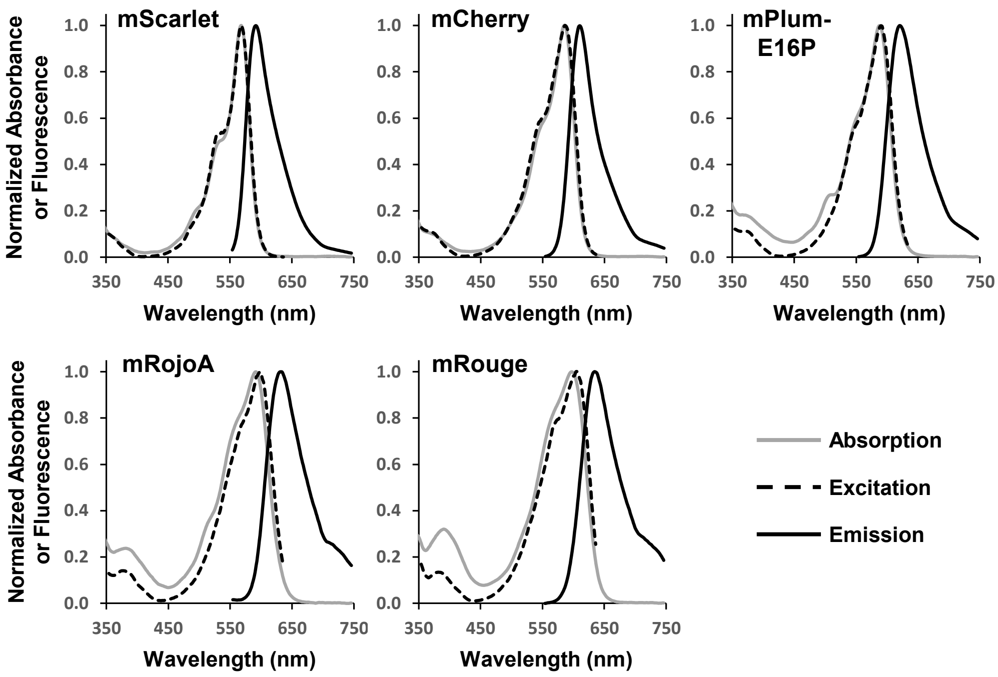


**Supplementary Figure 2. Spectral characterization of various RFPs.** Absorption, excitation ($\lambda_{em}$ = 655 nm) and emission ($\lambda_{ex}$ = 535 nm) spectra were measured in phosphate-buffered saline (137 mM sodium chloride, 2.7 mM potassium chloride, 10 mM disodium phosphate, 1.8 mM monopotassium phosphate, pH 7.4).

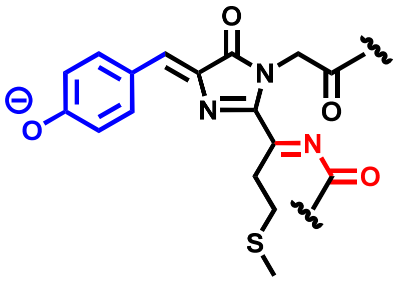

**Supplementary Figure 3. RFP chromophore structure.** The chromophore in mCherry and its variants is derived from a Met-Tyr-Gly tripeptide. The phenolate and acylimine groups are shown in blue and red, respectively.

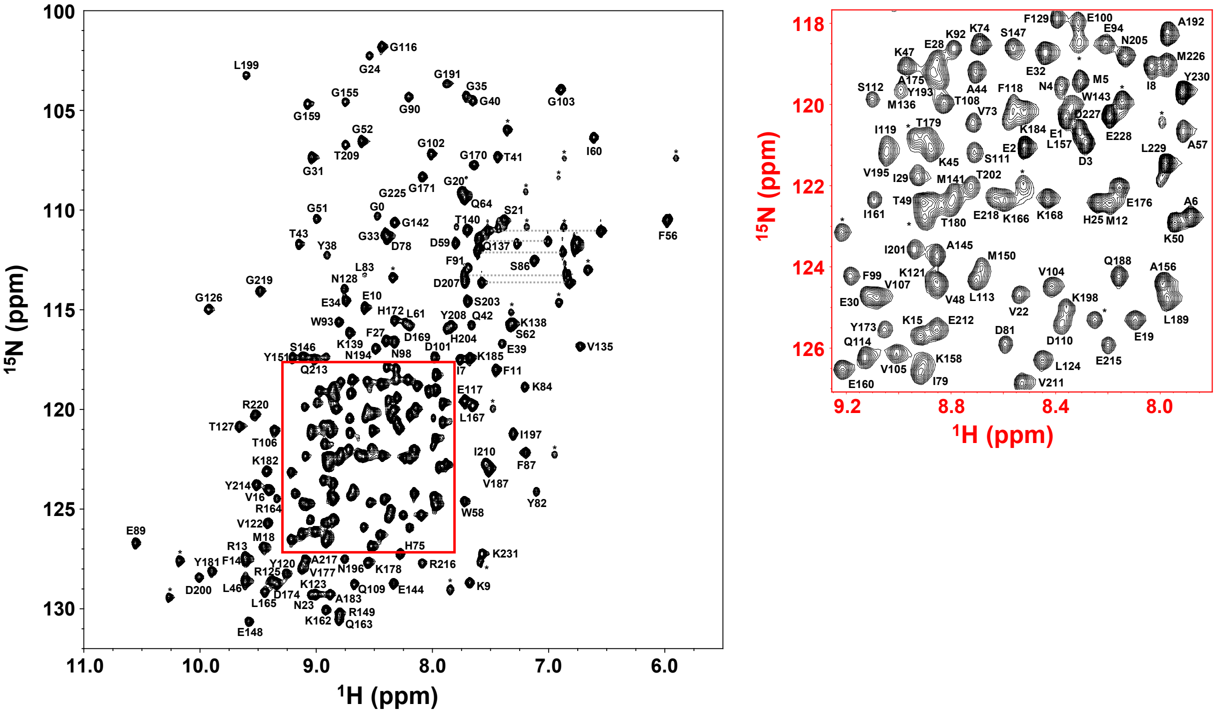


**Supplementary Figure 4. Assigned $^{1}$H-$^{15}$N HSQC spectrum of mCherry.** Side-chain amide resonances from asparagine and glutamine residues are connected by horizontal lines.

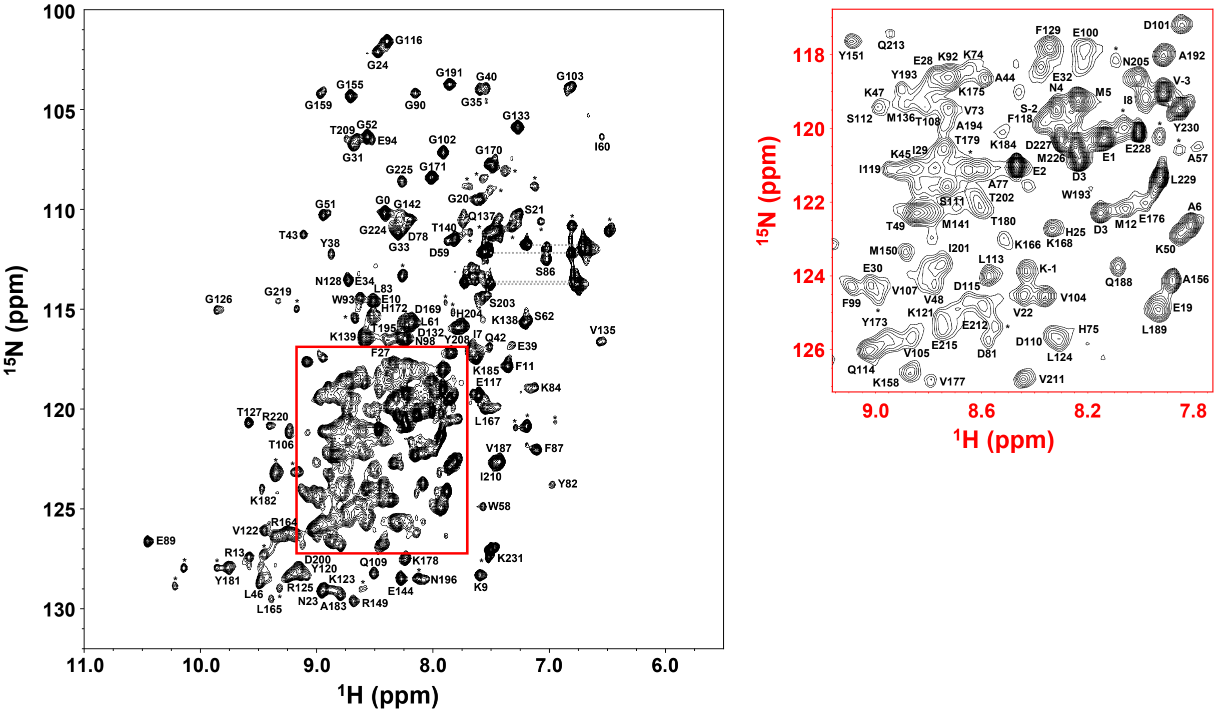


**Supplementary Figure 5. Assigned $^{1}$H-$^{15}$N HSQC spectrum of mRouge.** Side-chain amide resonances from asparagine and glutamine residues are connected by horizontal lines.

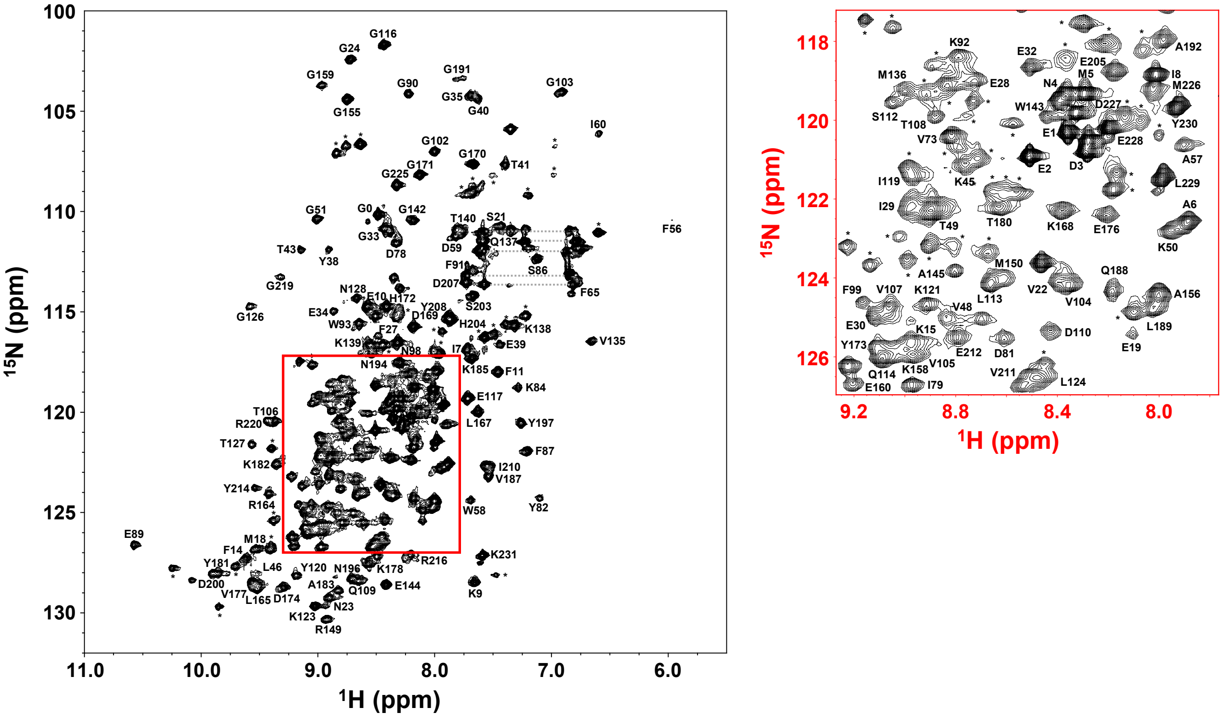


**Supplementary Figure 6. Assigned $^{1}$H-$^{15}$N HSQC spectrum of mRojoA.** Side-chain amide resonances from asparagine and glutamine residues are connected by horizontal lines.

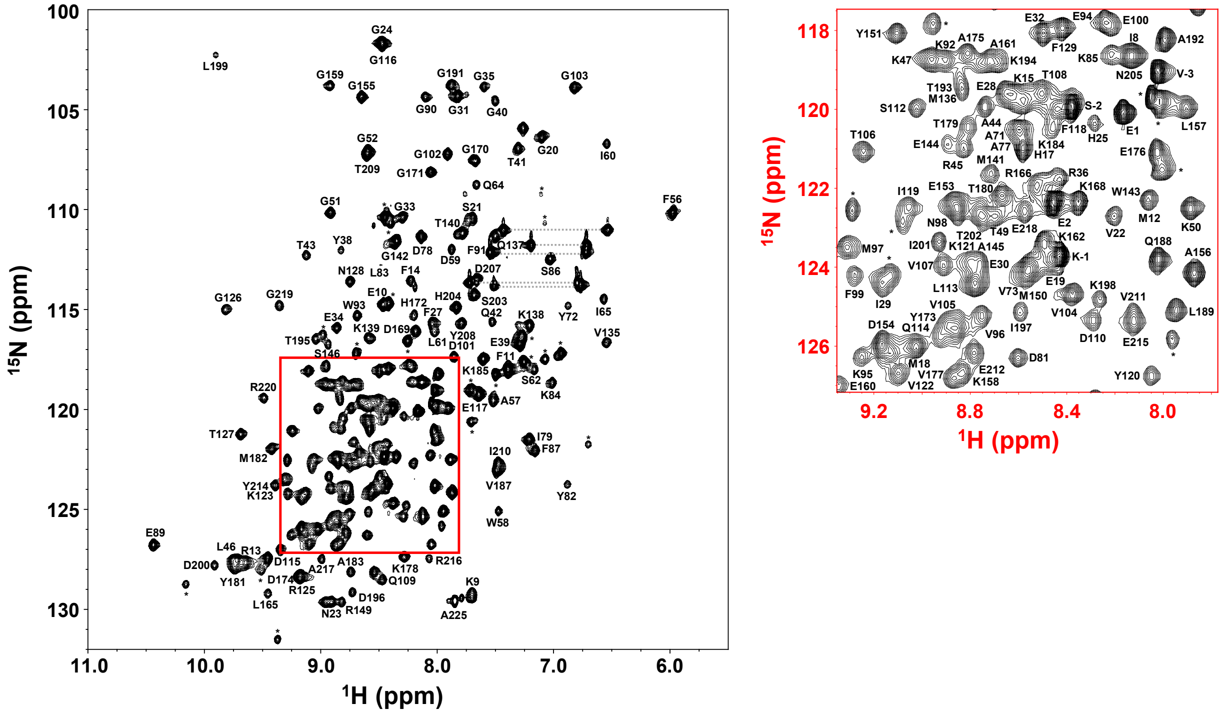


**Supplementary Figure 7. Assigned $^{1}H$-$^{15}N$ HSQC spectrum of mPlum-E16P.** Side-chain amide resonances from asparagine and glutamine residues are connected by horizontal lines.

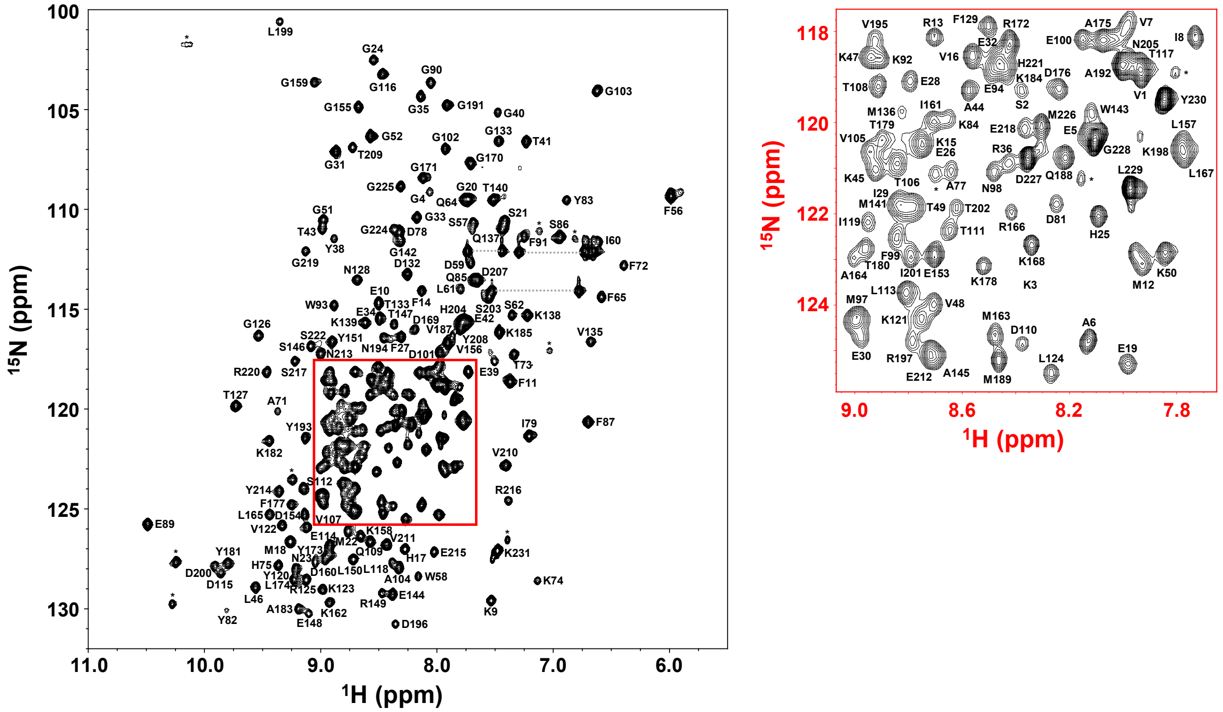


**Supplementary Figure 8. Assigned $^{1}$H-$^{15}$N HSQC spectrum of mScarlet.** Side-chain amide resonances from asparagine and glutamine residues are connected by horizontal lines.

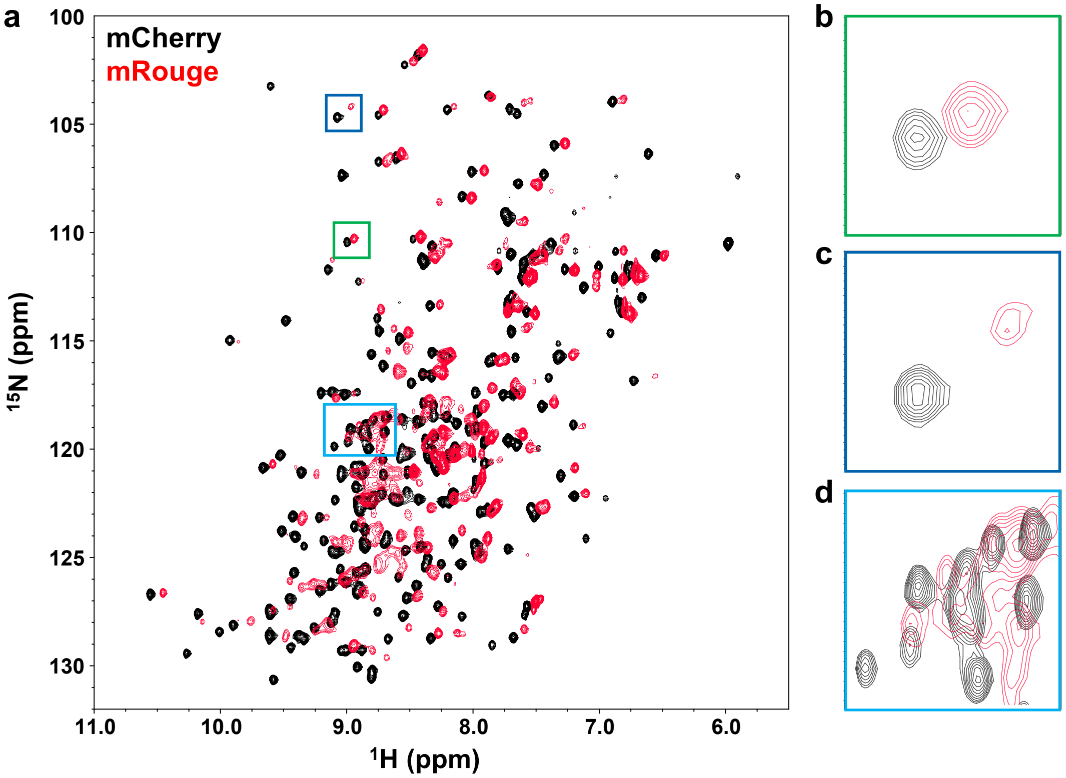


**Supplementary Figure 9. Dim RFPs exhibit peak broadening throughout their $^{1}$H-$^{15}$N HSQC spectrum.** (a) Overlaid HSQC spectra of the bright mCherry (black) and dim mRouge (red), normalized for concentration, show systematic peak broadening in mRouge. (b) Some rigid residues, such as G51 (shown), do not exhibit significant broadening in mRouge. (c) Other residues, such as G159 (shown), display substantial peak broadening and intensity loss compared to mCherry. (d) A clear loss of peak resolution due to broadening is observed in the central region of the HSQC spectrum (representative cut-out shown).

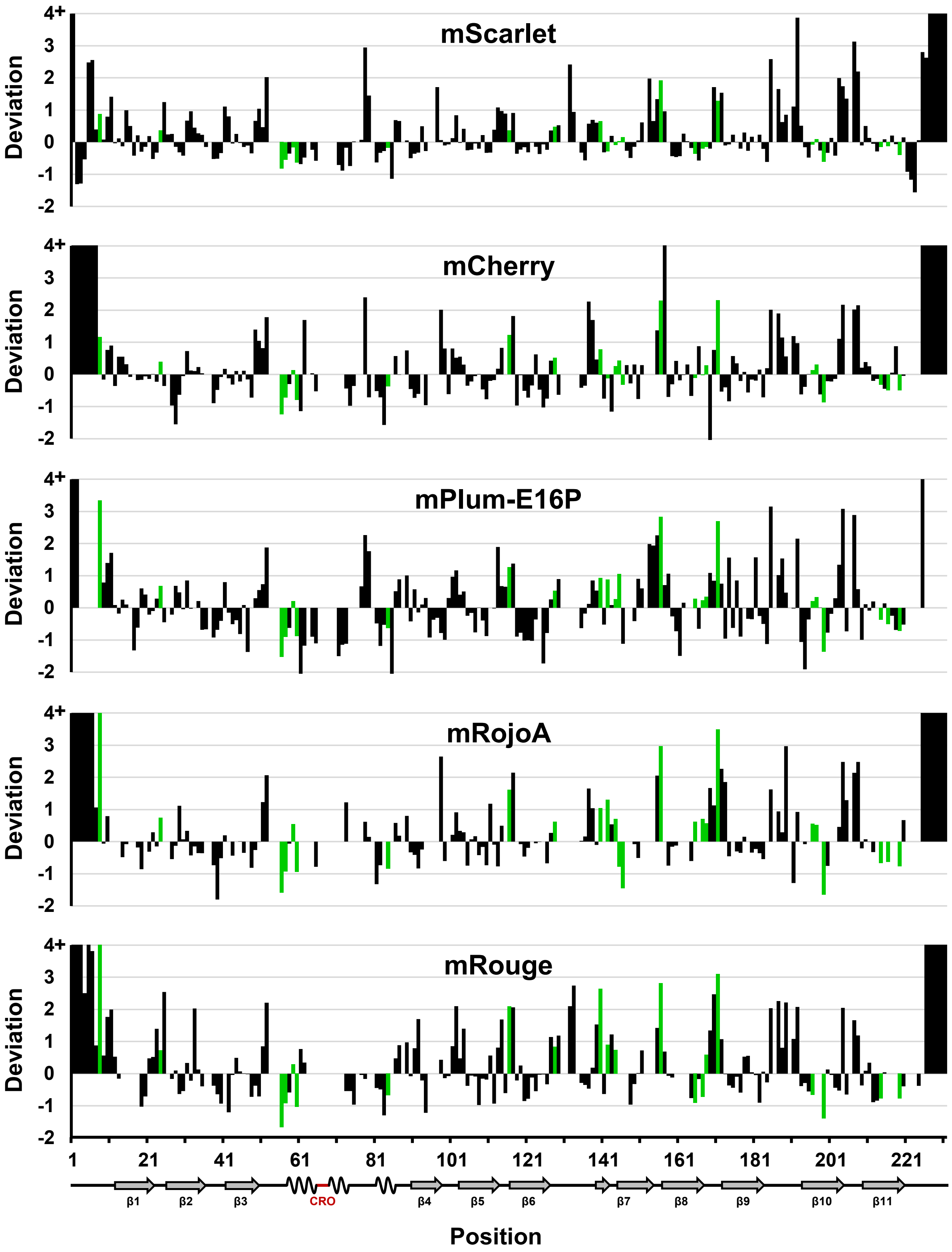


**Supplementary Figure 10. HSQC peak intensity deviation by residue.** Histograms show peak intensity deviations for each protein, with green bars indicating residues where a negative correlation between intensity deviation and quantum yield is observed, reflecting a positive correlation between rigidity and brightness. mCherry secondary structure elements are shown below the x-axis for reference.

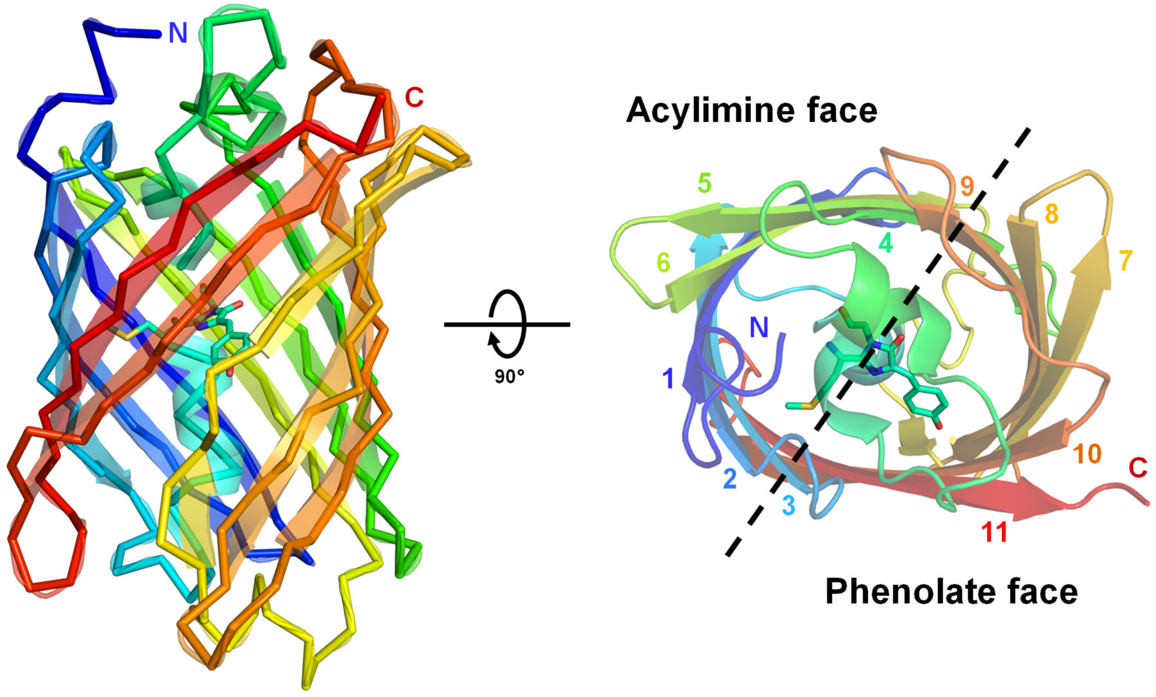


**Supplementary Figure 11. RFP β-strand ordering.** An archetypal RFP β-barrel structure from the mCherry crystal structure (PDB ID: 2H5Q) is shown colored by a blue-to-red rainbow spectrum from N-terminus to C-terminus to highlight strand ordering. On a top-down view of the barrel, strands are numbered accordingly. The phenolate and acylimine faces of the RFP barrel comprise beta strands 7–11 and 1–6, respectively.

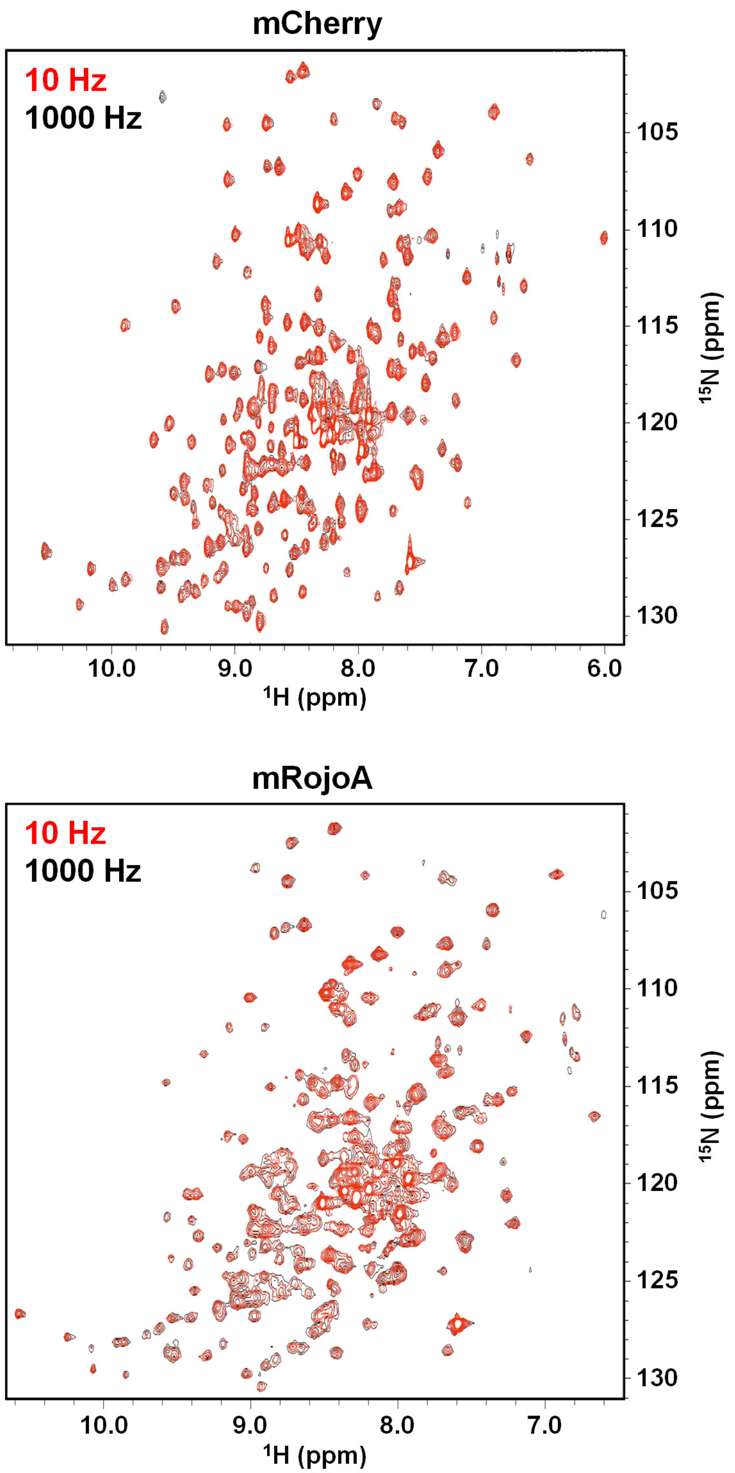


**Supplementary Figure 12. RFPs are insensitive to CPMG relaxation-dispersion experiments.** CPMG relaxation-dispersion spectra were collected at field strengths from 10 Hz to 1000 Hz for the bright mCherry and dim mRojoA RFPs. In both proteins, spectra at these extremes were nearly superimposable, with only minor differences in peak volumes and intensities for most residues. These results indicate that RFPs are rigid on the timescale and/or range of motions probed by CPMG.

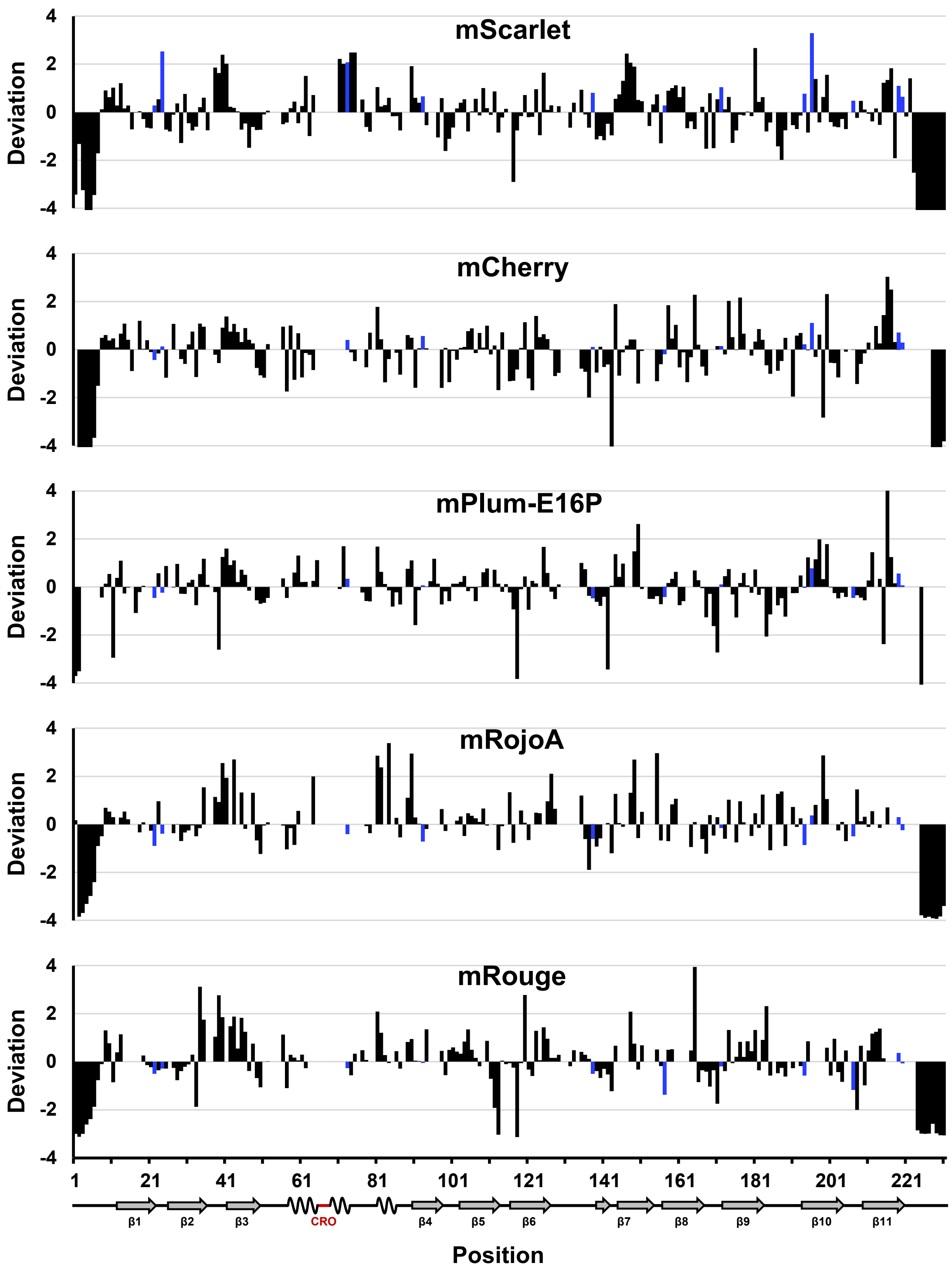


**Supplementary Figure 13. Normalized correlation time by residue.** Histograms show normalized correlation times for each protein, with blue bars marking positions where correlation time positively correlates with quantum yield (i.e., greater rigidity corresponds to higher quantum yield). Median correlation times were 16.3 ns, 16.5 ns, 14.2 ns, 18.4 ns, and 16.6 ns for mScarlet, mCherry, mPlum-E16P, mRojoA, and mRouge, respectively. mCherry secondary structure elements are shown along the x-axis.

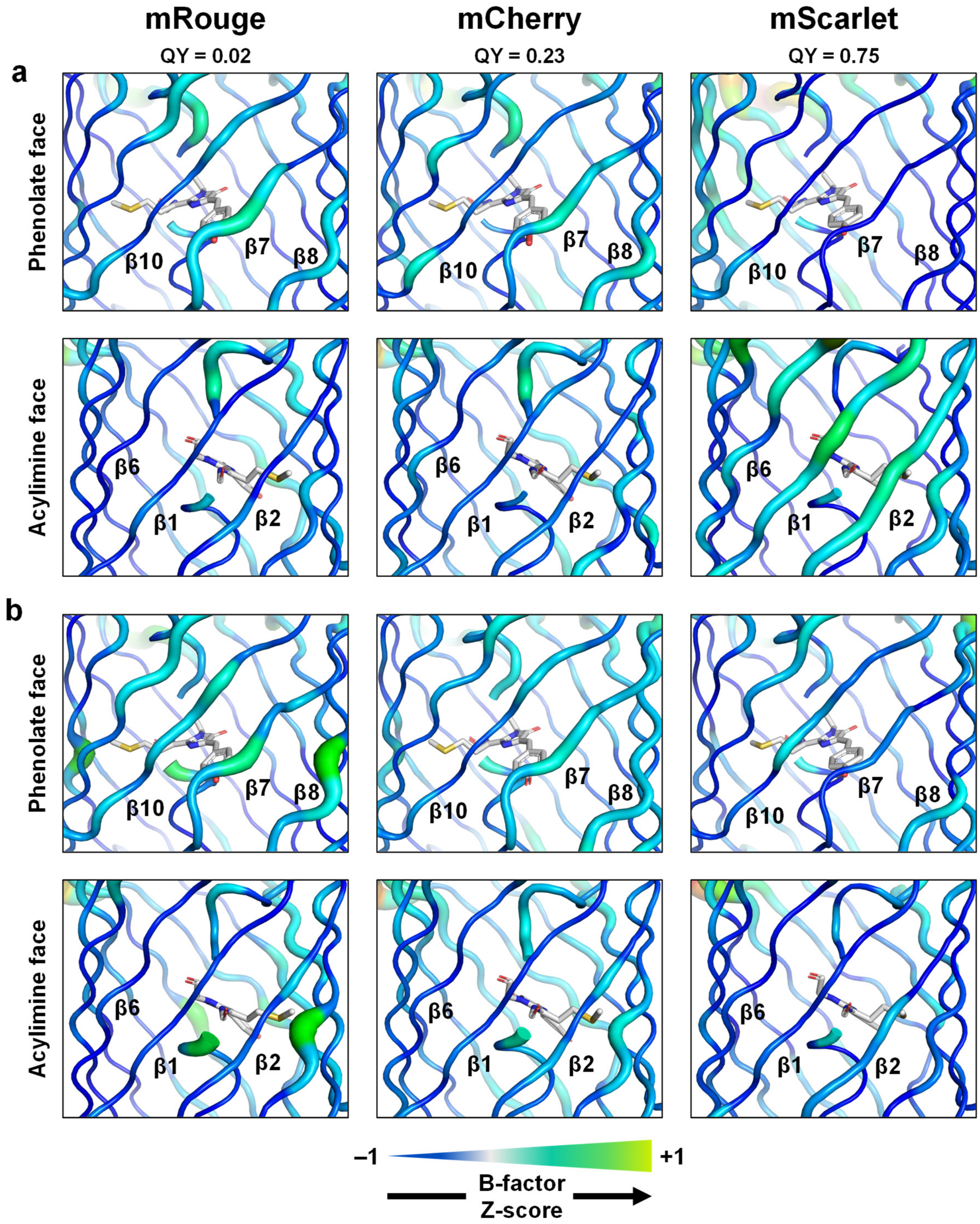


**Supplementary Figure 14. Conformational heterogeneity in RFP crystal structures.** B-factor Z-scores for each residue ($C_\alpha$) are represented as putty plots, with thickness proportional to the Z-score. Positive and negative Z-scores indicate increased flexibility or rigidity relative to the average residue in the protein, respectively. The chromophore is shown as white sticks. (a) At non-cryogenic temperature (277 K), β-strands β7, β8, and β10 on the phenolate face generally become more rigid as the quantum yield (QY) increases, whereas β-strands β1, β2, and β6 on the acylimine face are more flexible in mScarlet. (b) At cryogenic temperature (100 K), β-strands β7, β8, and β10 on the phenolate face become more rigid as QY increases, while β-strands β1, β2, and β6 on the acylimine face remain mostly unchanged.

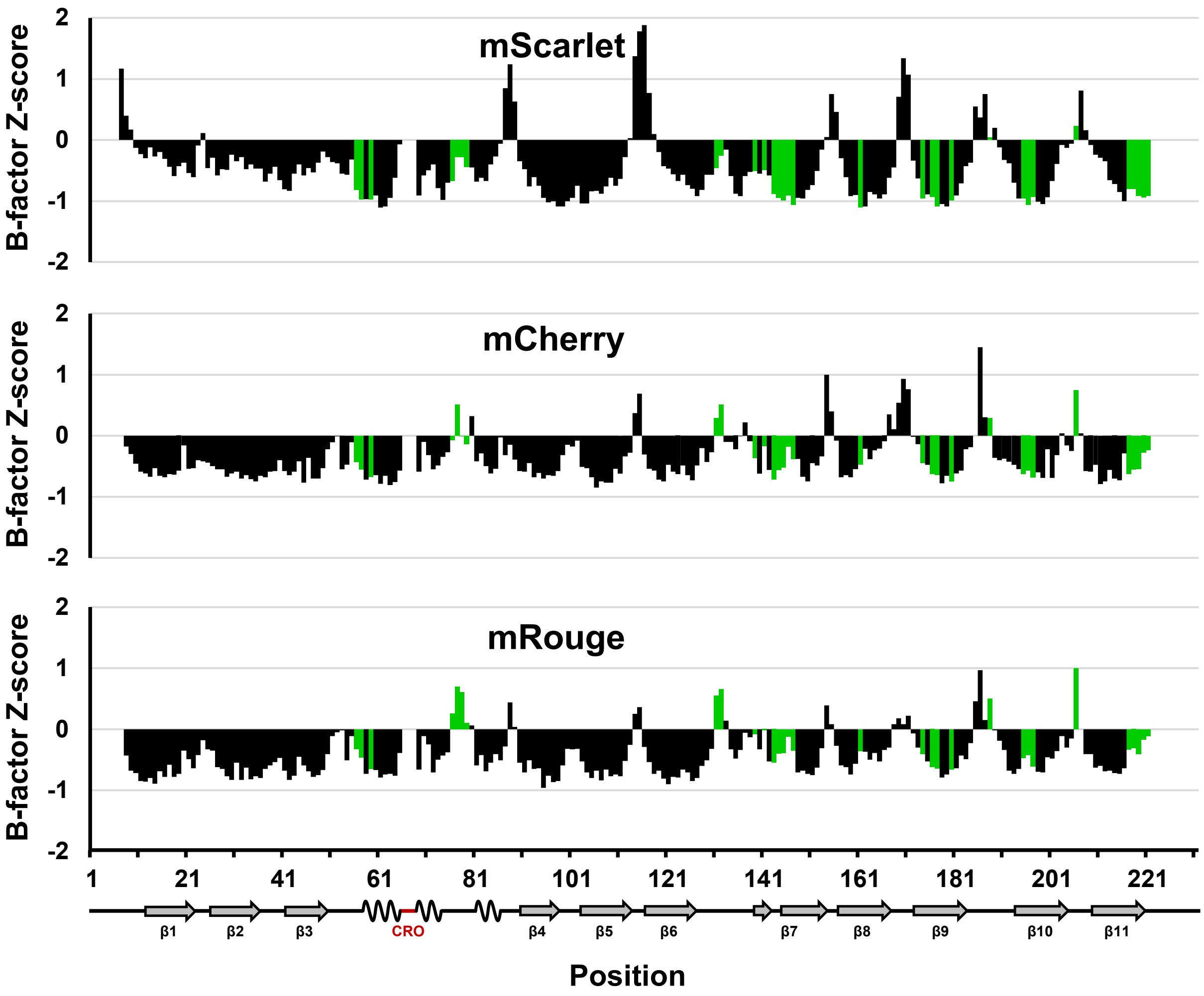


**Supplementary Figure 15. Non-cryogenic temperature (277 K) B-factor Z-scores by residue.** Histograms display the Cα B-factor Z-scores for every residue of each RFP. Green bars indicate residues exhibiting an inverse correlation between B-factor Z-score and quantum yield, suggesting that increased structural rigidity enhances fluorescence brightness. For reference, mCherry secondary structure elements are annotated below the x-axis.

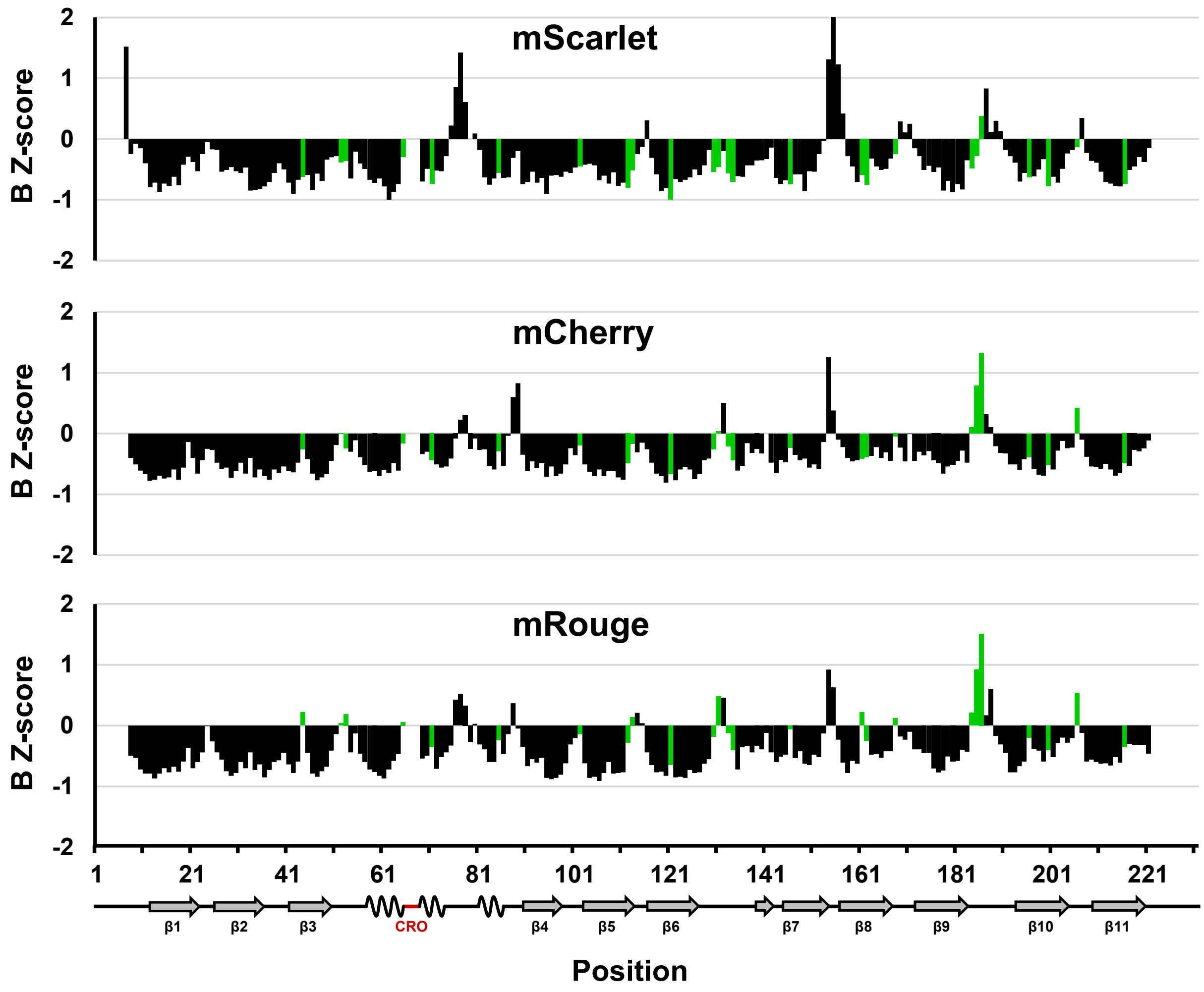


**Supplementary Figure 16. Cryogenic temperature (100 K) B-factor Z-scores by residue.** Histograms display the Cα B-factor Z-scores for every residue of each RFP. Green bars indicate residues exhibiting an inverse correlation between B-factor Z-score and quantum yield, suggesting that increased structural rigidity enhances fluorescence brightness. For reference, mCherry secondary structure elements are annotated below the x-axis.

**Positive Correlations**

**Exact Matches**

| | Total | Matches: NMR | X-ray, 277 K | X-ray, 100 K |
|---|---|---|---|---|
| **NMR** | 24 | | 9 | 4 |
| **X-ray, 277 K** | 31 | 9 | | 5 |
| **X-ray, 100 K** | 26 | 4 | 5 | |

**$C_\alpha$ Distance < 7 Å**

| | Total | Matches: NMR | X-ray, 277 K | X-ray, 100 K |
|---|---|---|---|---|
| **NMR** | 24 | | 17 | 14 |
| **X-ray, 277 K** | 31 | 22 | | 18 |
| **X-ray, 100 K** | 26 | 12 | 12 | |

**Negative Correlations**

**Exact Matches**

| | Total | Matches: NMR | X-ray, 277 K | X-ray, 100 K |
|---|---|---|---|---|
| **NMR** | 24 | | 7 | 4 |
| **X-ray, 277 K** | 30 | 7 | | 8 |
| **X-ray, 100 K** | 21 | 4 | 8 | |

**$C_\alpha$ Distance < 7 Å**

| | Total | Matches: NMR | X-ray, 277 K | X-ray, 100 K |
|---|---|---|---|---|
| **NMR** | 24 | | 18 | 15 |
| **X-ray, 277 K** | 30 | 20 | | 22 |
| **X-ray, 100 K** | 21 | 13 | 17 | |

**Supplementary Figure 17. Comparison of residue positions showing correlations between dynamics and quantum yield identified by NMR or X-ray crystallography.** Positive and negative correlations indicate residues that become more rigid or more flexible as quantum yield increases, respectively. Residue positions from each originating dataset (rows) are considered matched to a comparator dataset (columns) if a corresponding correlation is found at the same position (Exact Matches) or within a 7.0 Å $C_\alpha$–$C_\alpha$ distance ($C_\alpha$ Distance < 7.0 Å) to a correlated position in the comparator dataset. This distance cutoff was chosen to allow matches across adjacent residues on neighboring β-strands.

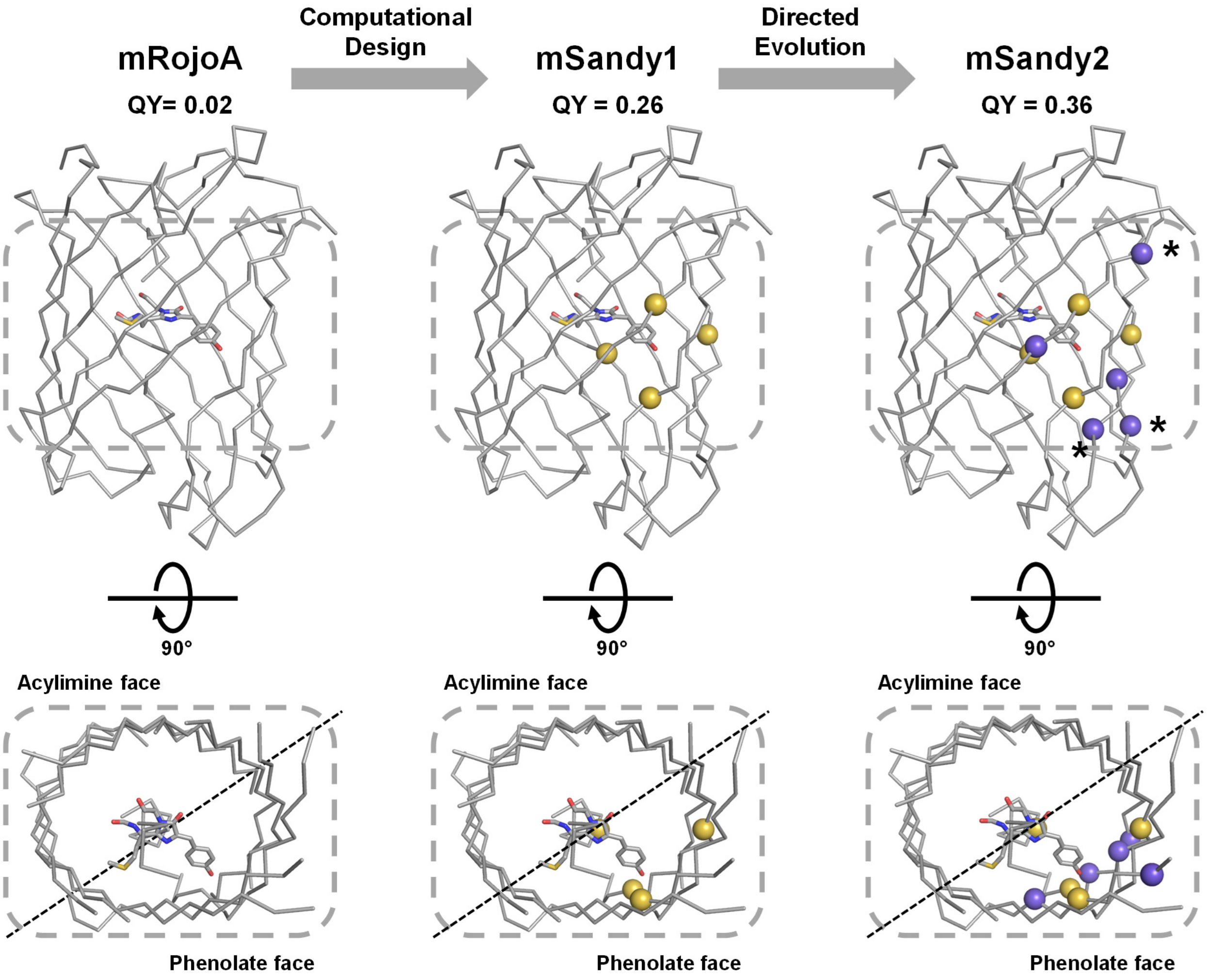


**Supplementary Figure 18. Engineering the bright RFP mSandy2 from dim mRojoA.** To increase the quantum yield (QY) of mRojoA, we used computational design to tighten packing around the chromophore phenolate [16], generating mSandy1. This introduced four first-shell mutations around the chromophore (yellow spheres)—three on the phenolate face (W143S, I161V, Y197I) and one on the central helix (P63L). mSandy1 also contains the T16V mutation on the acylimine face (not shown) that we introduced to facilitate chromophore maturation. Directed evolution then produced mSandy2, adding five more phenolate-face mutations (blue spheres, L163V, L165F, D174V, N194Y, L199M), three of which are located distal to the chromophore (asterisks). The chromophore is shown as sticks. In all cases, residues are mapped onto the mRojoA crystal structure (PDB ID: 3NEZ).
